# Liquid liquid phase separation of the intrinsically disordered protein JPT2 compartmentalizes components of NAADP-evoked Ca^2+^signaling

**DOI:** 10.64898/2026.01.10.698830

**Authors:** Sushil Kumar, Gihan S. Gunaratne, Jasmine Cornish, Kirsten M. Silvey, Qianru Mu, Zhong Guan, Gabriella T. Heller, James. T. Slama, Sandip Patel, Jonathan S. Marchant

**Affiliations:** Department of Cell Biology, Neurobiology & Anatomy, Medical College of Wisconsin, 8701 Watertown Plank Road, Milwaukee, WI 53226, USA; The Francis Crick Institute, London, NW1 1AT, UK; Bind Research, Apex, 1 Tribeca Walk, London, NW1 0QE; Structural and Molecular Biology, University College London, Gower Street, London, WC1E 6BT, UK; Department of Cell and Developmental Biology, University College London, Gower Street, London WC1E 6BT, UK; Department of Medicinal & Biological Chemistry, University of Toledo College of Pharmacy and Pharmaceutical Sciences, 3000 Arlington Avenue, Toledo, OH 43614, USA

## Abstract

Nicotinic acid adenine dinucleotide phosphate (NAADP) is a Ca^2+^-releasing second messenger that activates two-pore channels (TPCs) on endosomes and lysosomes. Rather than binding TPCs directly, NAADP acts through cytoplasmic NAADP-binding proteins (NAADP-BPs) which are essential for endolysosomal Ca^2+^ release. Here we characterized the properties of two recombinant, purified NAADP-BPs: Jupiter Microtubule Associated Homolog 2 (JPT2) and like-Sm protein 12 (LSM12). In contrast to LSM12, JPT2 is predicted to be an intrinsically disordered protein, a feature confirmed by circular dichroism and NMR spectroscopy. Under conditions of low Na^+^ concentration or molecular crowding, JPT2 underwent phase separation, as demonstrated by multiple orthogonal approaches. JPT2 condensates displayed liquid-like behavior and efficiently recruited LSM12, a novel fluorescent NAADP analog, as well as tubulin. JPT2 condensates also interacted with polymerized microtubules and lysosomes isolated from human cell lines. These findings reveal an unexpected capability of NAADP-BPs to undergo phase separation, and segregate with components needed for NAADP-dependent Ca^2+^ release. We speculate that these signaling condensates dictate cellular NAADP sensitivity, desensitization of NAADP responses, as well as NAADP targeting to TPCs at membrane contact sites between acidic organelles and the endoplasmic reticulum.

**Graphical Abstract:** 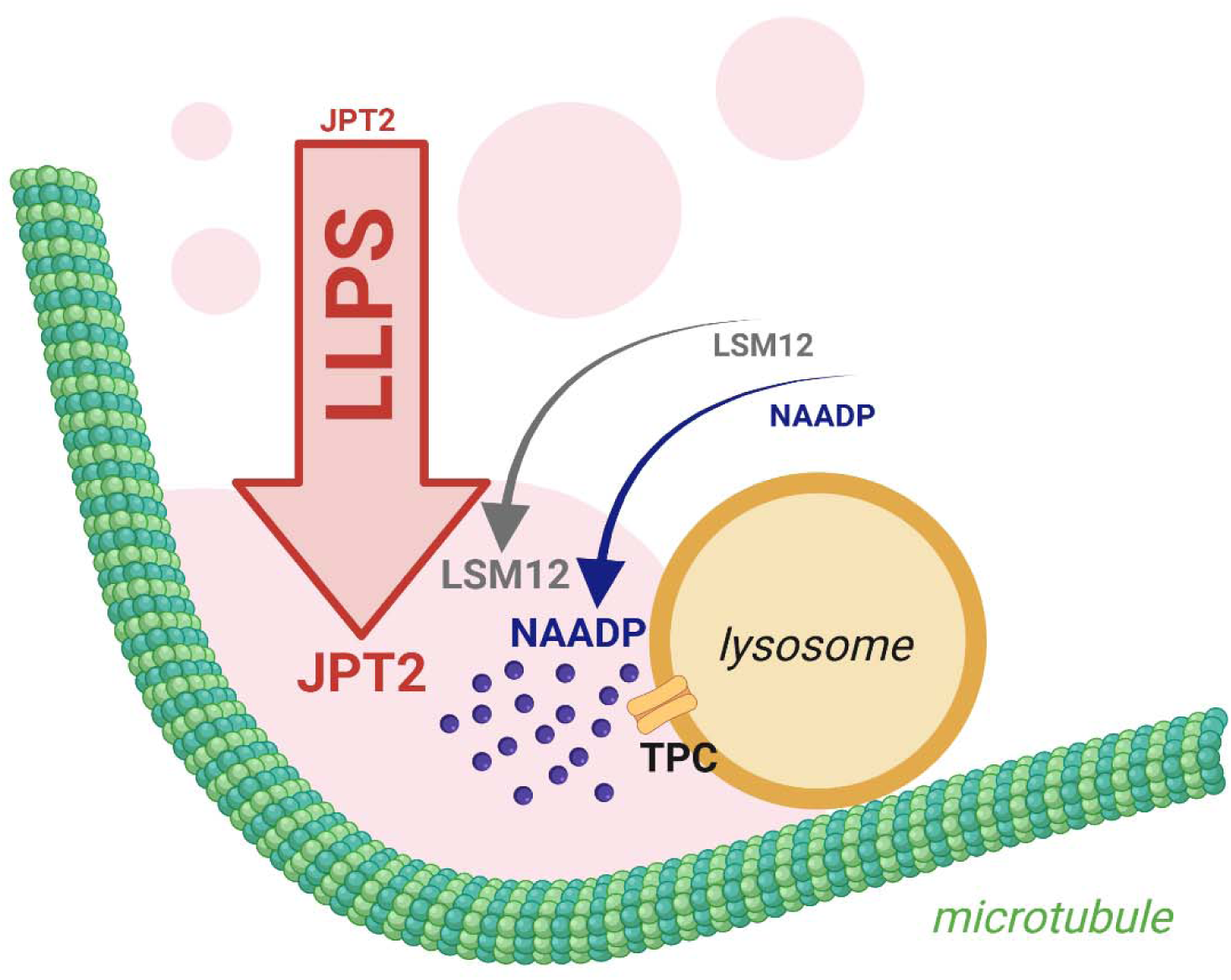

## Introduction

Nicotinic acid adenine dinucleotide phosphate (NAADP) is a nucleotide-based second messenger that mobilizes Ca^2+^ in many different cell types. NAADP-evoked Ca^2+^ signals regulate a wide range of physiological processes, and dysregulation of this pathway is implicated in a growing number of disease states [1–5].

The fundamental properties of NAADP-evoked Ca^2+^ signaling were defined decades ago, illuminated through studies of the robust Ca^2+^ response elicited by NAADP in sea urchin egg homogenates [6–8]. These studies revealed several distinctive features of the pathway: (i) an exceptional potency, with NAADP acting in the nanomolar range, (ii) a mechanism distinct from other intracellular Ca^2+^ release pathways in that NAADP targeted Ca^2+^ stores unique from the endoplasmic reticulum, and (iii) an apparent irreversibility of ^32^P-NAADP binding to its target underlying a phenomenon of self-inactivation whereby low NAADP concentrations desensitize Ca^2+^ release to subsequent additions of NAADP [9–13]. NAADP responsiveness is also capricious, with some mammalian cell types displaying robust responses to NAADP, while others appear unresponsive. These characteristics must ultimately relate to the properties and/or choreography between the molecular components of this Ca^2+^ signaling pathway.

Substantial progress has been made in defining the molecular basis of NAADP-evoked Ca^2+^ signaling. Key components include various enzymes responsible for NAADP synthesis and degradation [14–16], as well as the ion channels targeted by this second messenger. NAADP activates a family of endolysosomal ion channels known as two pore channels (TPCs), evolutionarily ancient members of the voltage-gated ion channel superfamily [17–20]. In human cells, two distinct family members – TPC1, TPC2 – are expressed. TPC1 expression is biased towards endosomes and TPC2 is predominantly localized within lysosomes. TPCs are permeable to both Na^+^ and Ca^2+^ and their ionic permeability is keyed to the mode of TPC activation [21, 22]. The endogenous phospholipid PI(3,5)P_2_ causes robust Na^+^ currents, whereas NAADP triggers Ca^2+^ release [23, 24]. Further, NAADP targets TPCs at sites of close membrane apposition between endolysosomal Ca^2+^ stores and the endoplasmic reticulum (ER), known as membrane contact sites (MCSs). Ca^2+^ released via NAADP-dependent activation of TPCs engages Ca^2+^-sensitive channels in the ER, thereby amplifying local signals into global cellular Ca^2+^ responses [25–27]. Therefore, by triggering Ca^2+^ signals at membrane contact sites, NAADP action is amplified beyond the local organellar neighborhood where TPCs reside.

Importantly, NAADP does not activate TPCs by binding directly to these ion channels. Instead, NAADP acts via accessory NAADP binding proteins (NAADP-BPs) that confer NAADP sensitivity to the channel complex [28–30]. Two such NAADP-BPs have been recently identified in mammalian cells – these are Jupiter Microtubule Associated Homolog 2 (JPT2, also known as HN1L [31, 32]) and like-Sm protein 12 (LSM12 [33]). Identification of these proteins provides new impetus to explain the fundamental properties of the NAADP signaling pathway [34, 35]. For example, it raises questions as to whether NAADP-BP expression correlates with manifestation of NAADP sensitivity, and whether their cellular localization dictates the spatial and temporal characteristics of NAADP action.

In this study, we examined the properties of the purified NAADP-BPs and demonstrate that JPT2 is an intrinsically disordered protein capable of undergoing liquid-liquid phase separation (LLPS) *in vitro*. JPT2 condensates selectively accumulate key components of the NAADP signaling pathway, including NAADP, LSM12, microtubules and lysosomes. The segregation of NAADP-BPs into liquid-like compartments distinct from the bulk cytoplasm offers a potential organizational framework for understanding several long-recognized features of NAADP-evoked Ca^2+^ signaling, including the variability in cell sensitivity to NAADP, response desensitization, and the spatial targeting of NAADP activity to membrane contact sites.

## Materials and Methods

### Materials and Reagents

Tobacco etch virus (TEV) protease was procured from GenScript. Isopropyl-β-D-thiogalactopyranoside (IPTG), lysozyme, polyethylene glycol 3350 (PEG3350), PEG silane (mPEG5K-silane), 1,6-hexanediol (1,6-HD), and the lysosomal isolation kit were sourced from Sigma. Polyacrylamide SDS-PAGE gels and Bio-safe Coomassie stain were sourced from Bio-Rad laboratories. Cell culture reagents, Alexa Fluor 488, Alexa Fluor 568, HisPur Ni-NTA columns, Zeba desalting and dye removal columns, and Lysotracker Deep Red were from Thermo Fisher Scientific. Fluorescently labeled Tubulin, GTP and taxol were obtained from Cytoskeleton. HEK293 cells were from American Type Culture Collection (ATCC). Protease inhibitor cocktail was from Roche. A plasmid encoding 6xHis-MBP-FUS was a gift from Nicolas Fawzi (Addgene plasmid #98651; http://n2t.net/addgene:98651; RRID:Addgene_98651). A bacterial expression construct containing an N-terminal 6xHis-tag fused upstream of a tobacco etch virus (TEV) protease recognition sequence and the coding sequence of human JPT2 (UniProt ID, Q9H910-3) was engineered within the pET28a vector. A similar bacterial expression construct containing an N-terminal 6xHis-GST tag, and TEV protease recognition sequence fused to LSM12 (UniProt ID, Q3MHD2) was synthesized by Genscript. Fluorescent NAADP analogs were synthesized by Zhong Guan and James Slama (University of Toledo, OH) from NADP as described in the Supplementary Methods.

### Bioinformatic analyses

For *in silico* analyses of cohorts of structured, disordered and liquid liquid phase separating (LLPS) proteins, the list of 1526 diverse fully folded proteins generated by Saar *et al.* [36] was used as the ‘structured’ dataset. The ‘disordered’ dataset was generated by retrieving accession codes for all proteins from the DisProt database [37] and filtering out entries annotated to have less than 50% disordered content. The ‘LLPS’ dataset was retrieved from PhaseSepDBv2.1 [38] and repeated entries were filtered out using accession codes. Unique accession codes from all lists were used to retrieve full length amino acid sequences from UniProtKB. The human proteome was also retrieved from UniProtKB. Primary sequence length, prevalence of individual amino acids, hydropathy values and net charge at physiological pH for all proteins were manually calculated in Microsoft Excel. Descriptive statistics were calculated in OriginLab. Differences in distribution of protein properties among datasets was tested for significance using the Mann-Whitney U-test. For radial plots, normalized values for protein hydropathy, fraction charged residue (FCR [39]), disorder (ESpritz [40]), Pi contacts (PiScore [41]), prion score (Prion Like Amino Acid Composition, PLAAC, [42]), LLPS propensity (catGRANULE [43]), cellular LLPS score (Proteins Involved in CoNdensates In Cells, PICNIC [44]), partner-driven phase-separation (PhaSepDB) and self-driving phase separation (PhaSepDB) [45]) were retrieved using the indicated bioinformatic tools. Prediction of Natural Disordered Regions (PONDR) scores were generated using PONDR VL-XT.

### Recombinant protein purification

For expression of recombinant proteins (JPT2, LSM12, FUS), individual plasmids were transformed into BL21 Star (DE3) *E. coli* cells (NEB) and a single colony was inoculated into 10 mL Luria Bertani (LB) broth with 100 mg/L ampicillin to be grown overnight in a gyratory incubator (37°C, 220 rpm). This starter culture was used to inoculate 1L of terrific broth (Sigma), with the culture incubated (37°C, 220 rpm) and then induced (at OD_600_ ∼1.2) with IPTG (0.5 mM) with further growth for 16-18 hours at 18°C. Cells were harvested by centrifugation and resuspended in ice-cold lysis buffer (50 mM Tris-HCl pH 8.0, 300 mM NaCl, 5% (v) glycerol, 0.1 mg/mL lysozyme, 5 mM imidazole) supplemented with EDTA-free protease inhibitor. The cell suspension was lysed by sonication and centrifuged (16,000 x g, 4°C for 1 hour). The supernatant was loaded on a HisPur Ni-NTA spin column and incubated on a nutator (4°C, 30 minutes). The column was washed (3 times) with 2 column volumes (CV) of wash buffer (50 mM Tris-HCl pH 8.0, 300 mM NaCl, 5% (v) glycerol, 10 mM imidazole), and the His-tagged proteins were eluted (3 times) with one CV of elution buffer (50 mM Tris-HCl pH 8.0, 300 mM NaCl, 5% (v) glycerol, 250 mM imidazole). Imidazole was removed by buffer exchange, and the N-terminal tags were removed using TEV protease. The solution was loaded on a Ni-NTA column, and the untagged proteins were collected in elution buffer containing 10 mM imidazole. The proteins were buffer exchanged in 25 mM HEPES, pH 7.4, 300 mM NaCl, and concentrated with 10 kDa cutoff Amicon filters (Millipore). For circular dichroism and NMR analyses, proteins were further purified by size-exclusion chromatography on a Superdex 75 column (GE Healthcare) at 5°C in the same buffer used for subsequent measurements. The final protein samples were flash frozen in liquid nitrogen in small aliquots, and stored at -80°C.

### Nuclear Magnetic Resonance (NMR) spectroscopy

^1^H-protein detected chemical shifts were performed on a 22.3 T Bruker AVANCE Neo spectrometer, equipped with a 5 mm ^1^H/^13^C/^15^N cryoprobe. ^1^H chemical shifts were referenced with respect to 4,4-dimethyl-4-silapentane-1-sulfonic acid (DSS). 1D ^1^H spectra were acquired with the standard zgesgp Bruker pulse sequence with 32,768 complex points, a spectral width of 14,700 Hz, and an acquisition time of 1.11 s. 16 scans were collected. Samples contained JPT2 at a concentration of 200 µM. Measurements were performed at 5 °C in 20 mM HEPES, 110 mM KCl, 10 mM NaCl, 1 mM TCEP, pH 7.2

### Circular Dichroism (CD)

CD spectra of 20 µM JPT2 were recorded on a Chirascan dichrograph (Applied Photophysics). Measurements were carried out at room temperature in a 0.1 mm path length quartz cuvette. Spectra were recorded in triplicate between 180 and 260 nm with a 0.5 nm increment and a 1 s integration time. Spectra were processed and baseline corrected using the Chirascan software. mDeg were converted to units of molar ellipticity per residue. All measurements were performed in 20 mM HEPES, 110 mM KCl, 10 mM NaCl, 1 mM TCEP, pH 7.2.

### NAADP-BP Protein labeling

Purified JPT2 and LSM12 were conjugated to either Alexa Fluor-488 or Alexa Fluor-568 (Invitrogen). Purified samples were buffer exchanged with sodium bicarbonate (0.1 M, pH 8.3) using Zeba™ spin desalting columns. Proteins were then mixed with the appropriate fluorophore at a 1:20 molar ratio and incubated for conjugation (1 hour at room temperature with continuous stirring). Unconjugated dye was removed using Zeba™ spin dye removal columns.

### Turbidity assays

Purified JPT2 protein (20 µM) was mixed with different volumes of ‘LLPS-0’ buffer (25 mM HEPES, pH 7.4) and ‘LLPS-300 buffer’ (25 mM HEPES, pH7.4, 300 mM NaCl) to obtain solutions of varied NaCl concentration. For the turbidity assay with 1,6-HD, JPT2 protein was diluted in LLPS-150 buffer (25 mM HEPES, pH7.4, 150 mM NaCl) with 10% PEG and supplemented with 1,6-HD to obtain indicated concentrations. Individual reaction conditions (100 µL) were set up in in triplicate in 1.5 mL tubes and immediately transferred to a 96-well plate (Costar). After incubation for 1 hour at room temperature, absorbance at 600 nm was measured using a SpectraMax i3x plate reader (Molecular Devices). Turbidity values were normalized to values recorded at the lowest NaCl or 1,6-HD concentration.

### Sedimentation assays

For sedimentation assays under crowding conditions, purified JPT2 (20µM) was incubated (25 mM HEPES 150 mM NaCl, pH 7.4) at room temperature for 30 minutes with various concentrations of polyethylene glycol (PEG3350). Samples were then centrifuged (21,000 *x g*, 4°C for 10 minutes). Supernatants were transferred to fresh tubes, and pellets were resuspended in the same volume of buffer as the supernatant. For sedimentation assays at varying salt concentrations, purified JPT2 (20µM) was incubated (room temperature, 30 minutes) in 25 mM HEPES buffer (pH 7.4) containing the indicated NaCl concentrations and then centrifuged (16,000 *x g*, 22°C for 30 minutes). Equal volumes of supernatant and pellet fractions were mixed with Laemmlii buffer (2x), heated (95°C, 5 minutes), and resolved by SDS-PAGE. Proteins were visualized by staining with Bio-safe Coomassie stain (Bio-Rad laboratories).

### Microscopy

JPT2 condensates were observed using differential phase contrast (DPC) or confocal microscopy using an Andor BC43 benchtop imaging system (Oxford Instruments, UK). Unless indicated otherwise, assays were set up in PEG silane coated, 18-chambered coverglass slides (#1.5 coverglass, Cellvis) to prevent surface-wetting. PEG-silane coating was performed as described in [46]. Images were captured with a 60x Plan Apochromat oil immersion objective. To observe the surface-wetting property of JPT2 droplets, assays were set up on non-PEG silane coated coverglass, and time-lapse imaging was performed with image acquisition every second. To prepare phase diagrams, the indicated concentrations of purified JPT2 protein were incubated with buffers containing the various concentrations of PEG3350 or NaCl and the presence or absence of droplets was scored. Images for a single experiment were captured using uniform acquisition settings.

### Fluorescence recovery after photobleaching (FRAP)

FRAP assays were carried out on a Nikon A1R laser scanning confocal microscope with a 60x Plan Apochromat oil immersion objective. Purified JPT2 protein (spiked with 1% Alexa Fluor-488 conjugated JPT2) was diluted into a low salt buffer (25 mM HEPES, pH 7.4, 25 mM NaCl) to a final concentration of 50 µM and immediately transferred to PEG-silane coated coverglass. Droplets of 2-3 µm diameter were selected, and a 0.3 µm-diameter circular region of interest (ROI) in the center of the droplet was photobleached by five 60 milliseconds laser pulses (λ=488nm). For quantitative comparison of fluorescence intensities, 0.3 µm diameter ROIs were selected on a non-photobleached droplet and within the bulk solution respectively and the mean fluorescence intensities of these ROIs were compared using NIS-Elements software. Images were acquired at 5s intervals with 14 images captured before photobleaching and 120 images after illumination. Measurements were then made for each time-point and processed using the following formula. Normalized intensity = (Intensity A– Intensity B) / (Intensity R – Intensity B), where A = mean intensity of the photobleached-ROI, B = mean intensity of the background-ROI, R = mean intensity of the reference-ROI. Fractional recovery was calculated by plotting the half-time of recovery to an exponential one-phase association curve (GraphPad Prism 10).

### Fluorescent NAADP assays

5-(AZ488-[CH_2_]_6_)-NAADP (abbreviated as AZ488-NAADP) and BODIPY-FL-(EG_4_)-NAADP (abbreviated as BODIPY-NAADP) were synthesized as reported in the Supplementary Methods. Partitioning of fluorescent NAADP into JPT2 condensates (50 µM JPT2, spiked with 1% AF568-JPT2) was monitored after inducing LLPS by incubating JPT2 in low-salt buffer (25 mM NaCl, 25 mM HEPES, pH 7.4) for 30 minutes, after which either fluorescent NAADP or the control unconjugated dye were added. Reactions were transferred onto PEG silane coated coverglass (18-chambered #1.5 coverglass, Cellvis) and images captured within 15-30 minutes of mixing. Fluorescent intensities inside and outside JPT2 condensates were measured using Fiji software. A partition coefficient ratio (enrichment) was calculated using the formula (mean fluorescence inside an individual droplet / mean fluorescence outside droplet).

### Biolayer interferometry

Biolayer interferometry (BLI) assays were carried out using an Octet BLI system (Sartorius). Purified recombinant JPT2 was biotinylated using EZ-Link Sulpho-NHS-LC-Biotin reagent (Thermo Fisher Scientific) and then buffer exchanged into an assay buffer containing 110 mM KCl, 10 mM NaCl, and 20 mM HEPES (pH 7.2) supplemented with 1% bovine serum albumin. Hydrated streptavidin (SAX) sensors (Sartorius) were equilibrated for 60 s in assay buffer followed by loading 50 nM of biotinylated JPT2 protein for 150 s. Next, a second equilibration step was performed for 60 s. Immobilized JPT2 protein was incubated with varying concentrations of AZ488-dye, AZ488-NAADP or BODIPY-NAADP for 180 s (association) followed by incubation with assay buffer for 180 s (dissociation).

### *In vitro* microtubule polymerization and co-incubation assays

*In vitro* polymerization assays utilized porcine brain tubulin tagged with HiLyte 488 (HL488) or HiLyte 647 (HL647) as a fluorescent label. Tubulin samples were resuspended in buffer (80 mM PIPES pH 6.9, 2 mM MgCl_2_, 0.5 mM EGTA) supplemented with 10% glycerol and 1 mM GTP to initiate polymerization. This mixture was incubated at 37°C for 20 minutes, and microtubules were stabilized by addition of taxol (20 µM). To study JPT2 association with microtubules, LLPS was induced by incubating JPT2 (50 µM, spiked with 1% AF568-JPT2) in low-salt buffer (25 mM HEPES pH 7.4, 25 mM NaCl, 30 minutes) followed by addition of taxol (20 µM) and GTP (1 mM), prior to combination (1:60 JPT2 sample) with the polymerized microtubule sample. Reactions were transferred onto PEG silane coated chambers (#1.5 coverglass, Cellvis) and images captured within 15-30 minutes of mixing.

### Lysosomal enrichment and coincubation assay

Lysosomes were isolated from HEK293 cells using a lysosome isolation kit (Sigma). Briefly, HEK293 cells were stained with the vital dye, LysoTracker Deep Red (50 nM) for 2 hours at 37°C. Cells were harvested by centrifugation (600 x g, 5 minutes), and washed with ice-cold PBS prior to another round of centrifugation (600 x g, 5 minutes at 4°C). The cell pellet was resuspended in an extraction buffer supplemented with a protease inhibitor cocktail for lysis using a Dounce homogenizer (20 strokes of Pestle B on ice). The lysed sample was centrifuged (1,000 x g, 4°C, 10 minutes) and the supernatant removed. This supernatant was again centrifuged (20,000 x g, 4°C, 20 minutes) and the resulting pellet resuspended in a minimal volume of extraction buffer (≤100µl) and stored on ice. This sample was used as the crude lysosomal fraction for imaging assays. To examine the relation between JPT2 condensates and lysosomes, LLPS was induced by incubating JPT2 (50 µM, spiked with 1% AF488-JPT2) in low-salt buffer (25 mM HEPES pH 7.4, 25 mM NaCl, 30 minutes) prior to addition of the lysosome preparation.

## Results

### *In silico* analysis of JPT2 properties

JPT2 is a small, basic protein containing four conserved PPGGxxSxxF motifs and a MASNIF motif shared across this gene family (**Fig. 1A**). Despite binding NAADP with high affinity [31, 35], JPT2 is predicted to be intrinsically disordered by both AlphaFold2 [47] and the disorder meta-predictor PONDR-FIT [48] (**Fig. 1B,C**). Consistent with these predictions, JPT2 is enriched in disorder-promoting amino acids (Lys, Pro, Glu, and Ser) and depleted of order-promoting residues (Cys, Ile, Trp, and Tyr) (**Fig. 1A** [49]). In contrast, LSM12 is predicted to be a structured NAADP-BP displaying a defined folded architecture (**Fig. 1B,C**, [50]).

**Figure 1.**
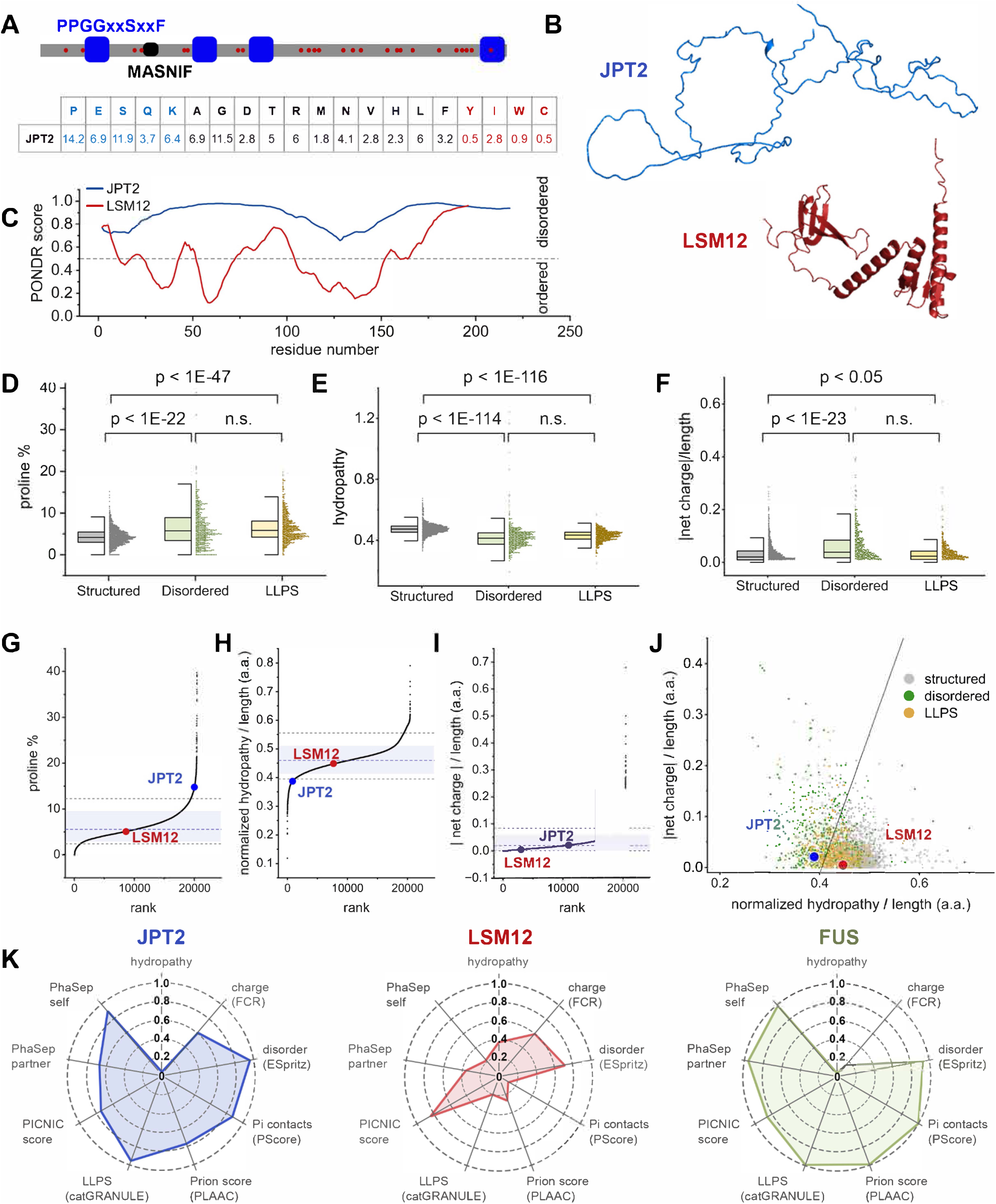
Bioinformatic analysis of JPT2 properties. **(A)** *Top,* schematic of JPT2 primary sequence, highlighting repeated PPGGxxSxxF motifs (blue), basic residues (red), and the MASNIF motif. *Bottom,* amino acid composition of JPT2. **(B)** Cartoon representations of AlphaFold2 structural predictions of JPT2 (blue, top) and LSM12 (red, bottom). **(C)** PONDR VL-XT prediction of IDRs in JPT2 (blue) and LSM12 (red); regions with scores >0.5 (dashed line) are classified as disordered. **(D-F)** Comparison of proline abundance (D), normalized hydropathy (E), and absolute net charge (F) among experimentally validated cohorts of fully structured proteins (grey, n=1562), proteins comprised of ≥ 50% IDRs (green, n=487), and LLPS-associated proteins (yellow, n=590). **(G-I)** Rank-ordered distributions of proline abundance (G), normalized hydropathy (H) and absolute net charge (I) across the human proteome. Dashed lines indicate median values, shaded regions represent standard error, and dotted lines denote the 5-95% confidence intervals. **(J)** scatter plot of absolute net charge versus normalized hydropathy for structured, disordered and LLPS-associated protein cohorts. The black line represents a reference boundary commonly used to distinguish ordered and disordered proteins. **(K)** Radial plots summarizing scores from nine predictive tools, ranked 0-1.0 across the human proteome. The LLPS-prone RNA-binding protein FUS (green) is shown as a reference. Hydropathy and fraction of charged residues (FCR) were calculated using CIDER; disorder was predicted with ESpritz; pi-pi contacts using PScore; prion-like amino acid composition using PLAAC; LLPS propensity using catGRANULE; condensate association using Proteins Involved in CoNdensates In Cells (PICNIC); and partner-driven versus self-driven phase separation using PhaSepPred,

Given the established role of intrinsic disorder in promoting liquid-liquid phase separation (LLPS), we compared the sequence properties of JPT2 with experimentally validated cohorts of fully structured proteins (n=1562), intrinsically disordered proteins (IDPs, n=487), and LLPS-associated proteins (n=590) (**Fig. 1D-F)**, [36, 38, 37]. Three parameters commonly associated with disordered proteins and LLPS propensity were examined: proline abundance, hydropathy and net charge. Median proline content was significantly higher in both disordered proteins (5.8%) and LLPS-associated proteins (5.8%) compared with structured proteins (4.2%) (**Fig. 1D**). Hydropathy, which is inversely correlated with LLPS propensity, was lower in disordered and LLPS-associated cohorts than in structured proteins (**Fig. 1E**). In addition, the absolute net charge was significantly higher in disordered and LLPS-associated proteins, consistent with a role for electrostatic interactions in LLPS (**Fig. 1F**) [51–54].

To place JPT2 and LSM12 within context of the human proteome, we rank-ordered these same parameters across all human proteins. JPT2 exhibited proline abundance and hydropathy values outside the 5-95% confidence intervals of the human proteome (**Fig. 1G,H**), whereas its net charge fell within the standard deviation (**Fig. 1I**). By contrast, LSM12 displayed near-median values for all three metrics. When absolute net charge was plotted against normalized hydropathy,

JPT2 clustered with the majority (>65%) of experimentally validated disordered proteins, while LSM12 clustered with structured proteins (92%) (**Fig. 1J**) [55].

Finally, we assessed LLPS propensity using various predictive tools. JPT2 scored highly for disorder (ESpritz), pi–pi contacts (PScore), prion-like amino acid composition (PLAAC), and LLPS propensity (catGRANULE), whereas LSM12 scored low across these metrics (**Fig. 1K**) [40, 42, 43, 39, 41]. Both proteins were predicted by PICNIC to localize to biomolecular condensates in cells, suggesting that LSM12 may partition into pre-existing condensates rather than drive their formation [45, 44]. Consistent with this, PhaSePred predicted JPT2 to preferentially undergo self-driven phase separation (score 0.90) rather than partner-dependent LLPS (0.68), while LSM12 scored weakly in both these categories [45, 44]. For comparison, the LLPS-prone RNA-binding protein FUS displayed a predictive profile more similar to JPT2 than to LSM12 (**Fig. 1K**) [56]. Collectively, this suite of bioinformatic analyses predict JPT2 as a disordered protein with likelihood to undergo LLPS in cells.

### Recombinant JPT2 is intrinsically disordered in solution

To enable biochemical and biophysical characterization, JPT2 was expressed recombinantly in *E. coli* and purified as described in Materials and Methods. To experimentally assess the conformational properties of recombinant JPT2, we performed circular dichroism (CD) spectroscopy and one-dimensional ^1^H NMR analysis under native aqueous conditions. The far-UV CD spectrum of JPT2 exhibited a pronounced minimum near ∼200 nm, with no detectable minima at 208 or 222 nm (**Fig. 2A**), indicating an absence of stable α-helical or β-sheet secondary structure. This spectral profile is characteristic of IDPs [57]. Consistent with these findings, the ^1^H 1D NMR spectrum of JPT2 displayed limited chemical shift dispersion in the amide proton region, with resonances clustered between approximately 6.0 and 8.5 ppm (**Fig. 2B**). The lack of well-dispersed amide resonances reflects conformational averaging and supports a predominantly disordered ensemble in solution [58].

**Figure 2.**
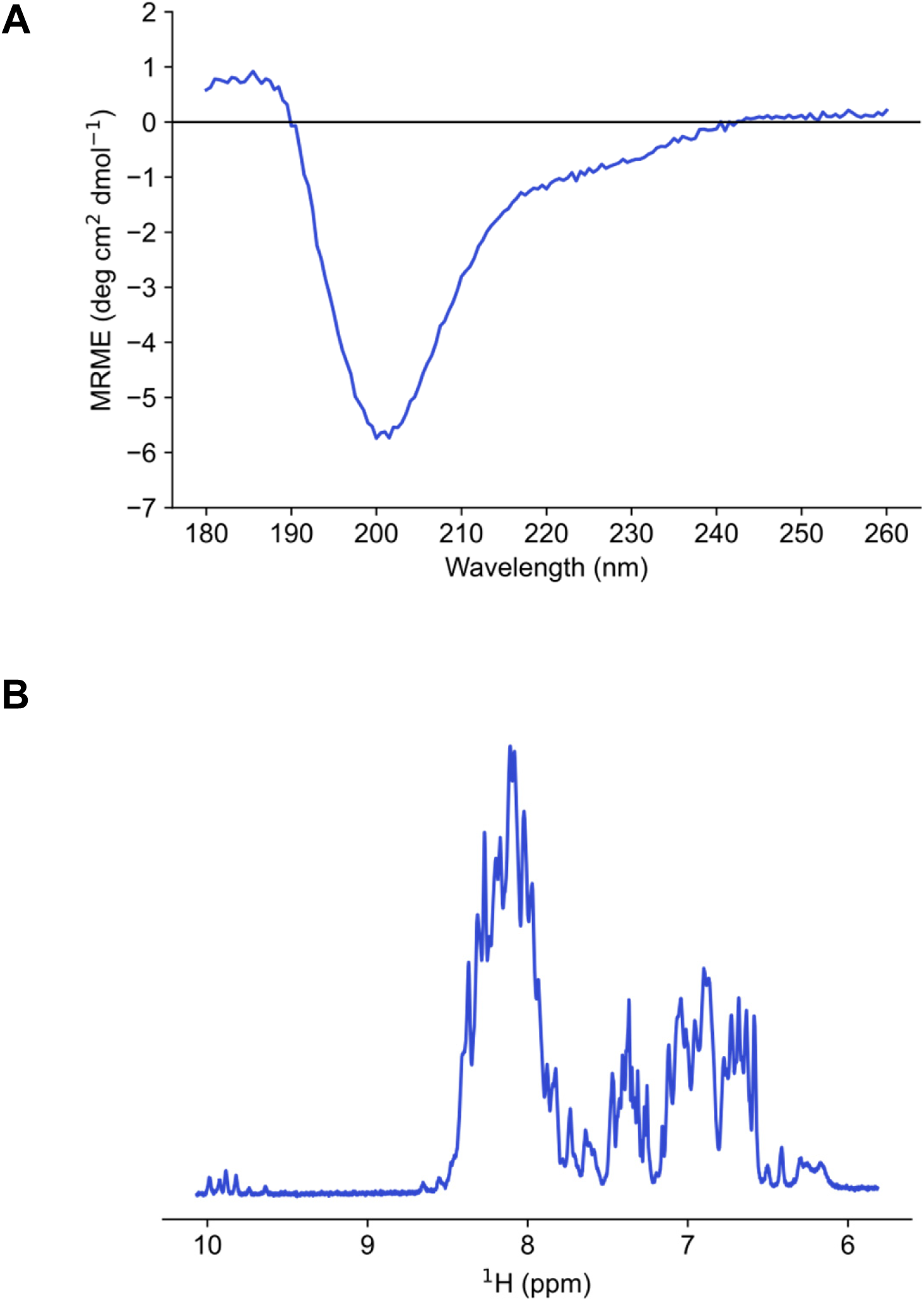
Biophysical experiments demonstrate that recombinant JPT2 is an intrinsically disordered protein. **(A)** Circular dichroism data of 20 µM JPT2.3 shows a single characteristic dip near 200 nm, known to correlate with disordered conformations [57]. **(B)** ^1^H 1D NMR spectrum of 200 µM JPT2 shows ^1^H chemical shift dispersion of amide regions between 6.0 - 8.5 ppm, characteristic of intrinsically disordered proteins [58].

### JPT2 undergoes LLPS *in vitro*

During purification of recombinant His-tagged JPT2, we noted that the protein solution was clear following the final affinity elution step but became rapidly turbid when diluted into TEV-cleavage buffer (**Fig. 3A**). This change coincided with a substantial reduction in ionic strength, from 300 mM NaCl in the elution buffer to 30 mM NaCl in the TEV cleavage buffer. Increasing the NaCl concentration back to 300 mM resulted in rapid clearing (Fig. 3A), indicating that the process was reversible and consistent with salt-dependent phase separation rather than irreversible aggregation. In line with this observation, we observed a decrease in quantitative turbidity measurements of JPT2 at higher NaCl concentrations (**Fig. S1**). For comparison, the LLPS-prone RNA-binding protein FUS [59] was also analyzed. Recombinant MBP-FUS became turbid following TEV treatment (**Fig. 3B**). In contrast to JPT2, however, increasing the NaCl concentration to 300 mM did not reverse FUS turbidity (**Fig. 3B**), suggesting that the molecular forces driving condensation differed between these proteins.

**Figure 3.**
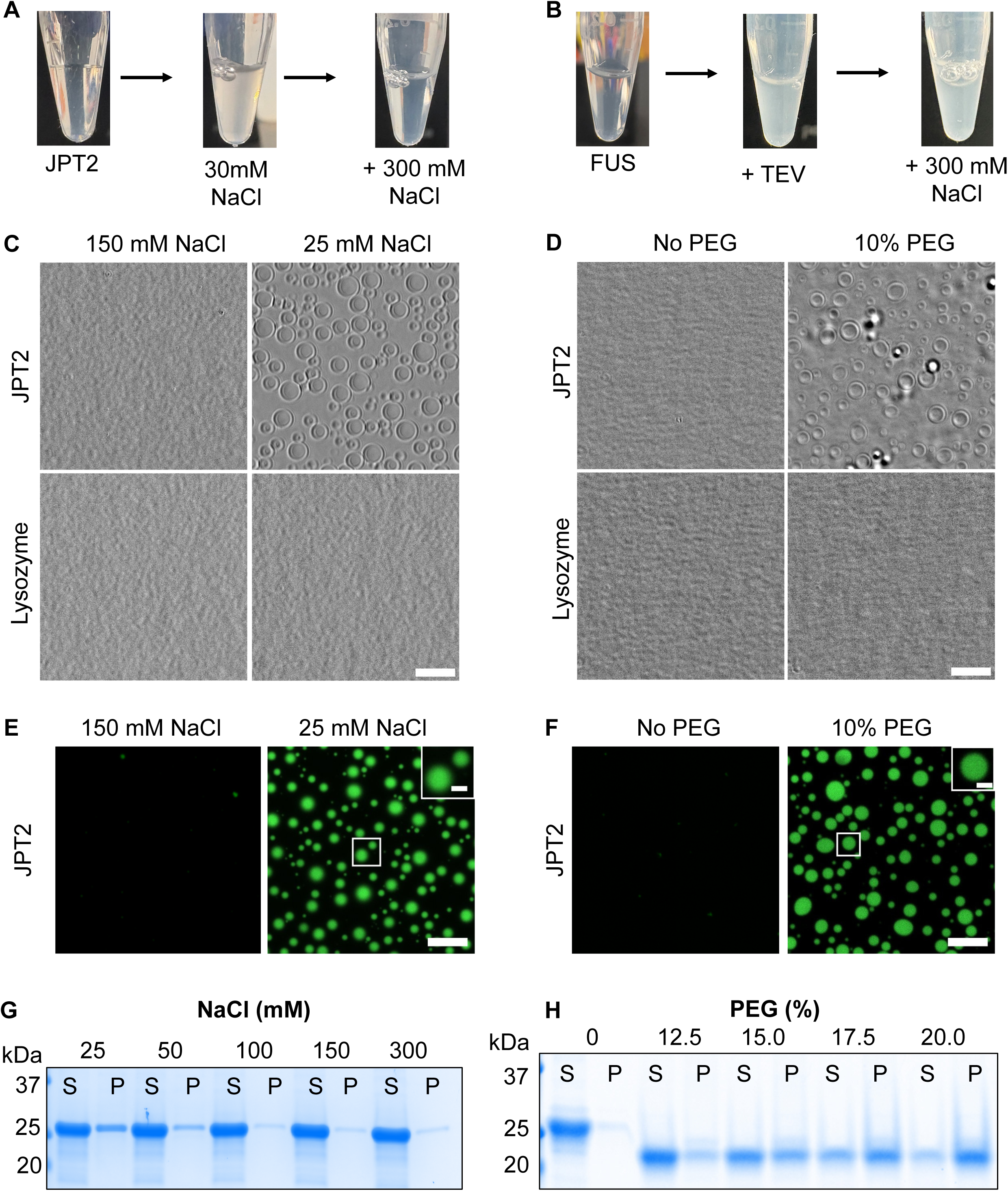
JPT2 undergoes phase separation in vitro. **(A)** A clear solution of recombinant His-tagged JPT2 in high-salt buffer (*left*) becomes turbid upon transfer to the low salt conditions in the TEV cleavage buffer (*middle*) and reverts to a clear solution upon supplementation with 300 mM NaCl (*right*). **(B)** A solution of recombinant MBP-FUS protein without TEV protease (*left*) becomes turbid following overnight incubation with TEV protease at 4°C (*middle*) and remains turbid after supplementation with 300 mM NaCl (*right*). **(C)** Representative differential phase contrast (DPC) images of JPT2 (50 µM; upper panels) or lysozyme (50 µM; lower panels) incubated for 1 hr at the indicated NaCl concentrations. **(D)** Representative DPC images of JPT2 (50 µM, upper panels) or lysozyme (50 µM, lower panels) incubated for 1 hour in the presence or absence of 10% PEG3350. **(E)** Representative confocal images of purified JPT2 (50 µM) incubated for 1 hour at the indicated salt concentrations. **(F)** Representative confocal images of purified JPT2 (50 µM) incubated for 1 hour in the presence or absence of 10% PEG3350. In both panels, unlabeled JPT2 was spiked with 1% Alexa Fluor 488-conjugated JPT2. Inset, enlarged views of individual JPT2 droplets. **(G, H)** Coomassie stained SDS-PAGE gels showing partitioning of purified JPT2 protein (20 µM) following incubation for 1 hour at the indicated NaCl (E) or PEG3350 (F) concentrations and subsequent centrifugation. P-pellet, S-supernatant. The size change of JPT2 in the presence of PEG is likely due to the formation of SDS-PEG micelles [99]. Scale bars: 10 µm (C, D, E, F), 2 µm (insets E, F)

We next examined JPT2 solutions using differential phase contrast (DPC) microscopy. At a low ionic strength (25 mM NaCl), JPT2 formed distinct spherical droplets of varying sizes, consistent with the formation of condensates, whereas solutions remained homogenous at higher salt concentrations (150 mM NaCl, **Fig. 3C**). This behavior was not observed for the control protein lysozyme under identical conditions (**Fig. 3C**). JPT2 condensation was also induced at high salt by the addition of the molecular crowding agent polyethylene glycol (10% PEG3350), while lysozyme again remained soluble under these conditions (**Fig. 3D**).

To directly visualize condensates, JPT2 was fluorescently labelled with Alexa Fluor-488 and mixed with unlabeled protein a 1:100 molar ratio. Confocal microscopy revealed spherical droplets under low salt conditions (**Fig. 3E**), with higher-magnification images showing relatively homogenous internal fluorescence within the droplet (**Fig. 3E, inset**). Similar droplets formed under PEG-induced crowding (**Fig. 3F**). Droplet sizes were comparable under low-salt and crowding conditions (mean areas of 5.3 ± 4.4 µm^2^ and 4.5 ± 4.0 µm^2^ respectively; **Fig. S2**). Therefore JPT2 undergoes autonomous phase separation driven by ionic strength and local protein concentration.

Finally, sedimentation assays provide an orthogonal measure of condensation. Decreasing NaCl concentration (**Fig. 3G**) or increasing PEG3350 concentration (**Fig. 3H**) resulted in progressive redistribution of JPT2 from the supernatant to the pellet fraction, consistent with phase separation observed using microscopy.

### Electrostatic interactions predominantly drive LLPS of JPT2

To define conditions governing JPT2 phase separation, phase diagrams were generated by titrating JPT2 concentration against either PEG3350 concentration or ionic strength. Results were plotted in ‘x-y’ format and summarized as a color-coded matrix. In the absence of PEG (150 mM NaCl), LLPS of JPT2 was not observed at low protein concentrations (≤20µM JPT2), and irregular assemblies formed at higher JPT2 concentrations (≥50µM JPT2) (**Fig. 4A**). Addition of PEG (10%) promoted formation of spherical droplets at substantially lower JPT2 concentrations (≥5 µM), with droplet number and size increasing with protein concentration (**Fig. 4A**). In the absence of crowding agents, JPT2 condensation was strongly salt-dependent, with droplets forming at low ionic strength and progressively dissolving as NaCl concentration increased (**Fig. 4B**). These findings suggest a prominent role for electrostatic interactions in JPT2 phase separation.

**Figure 4.**
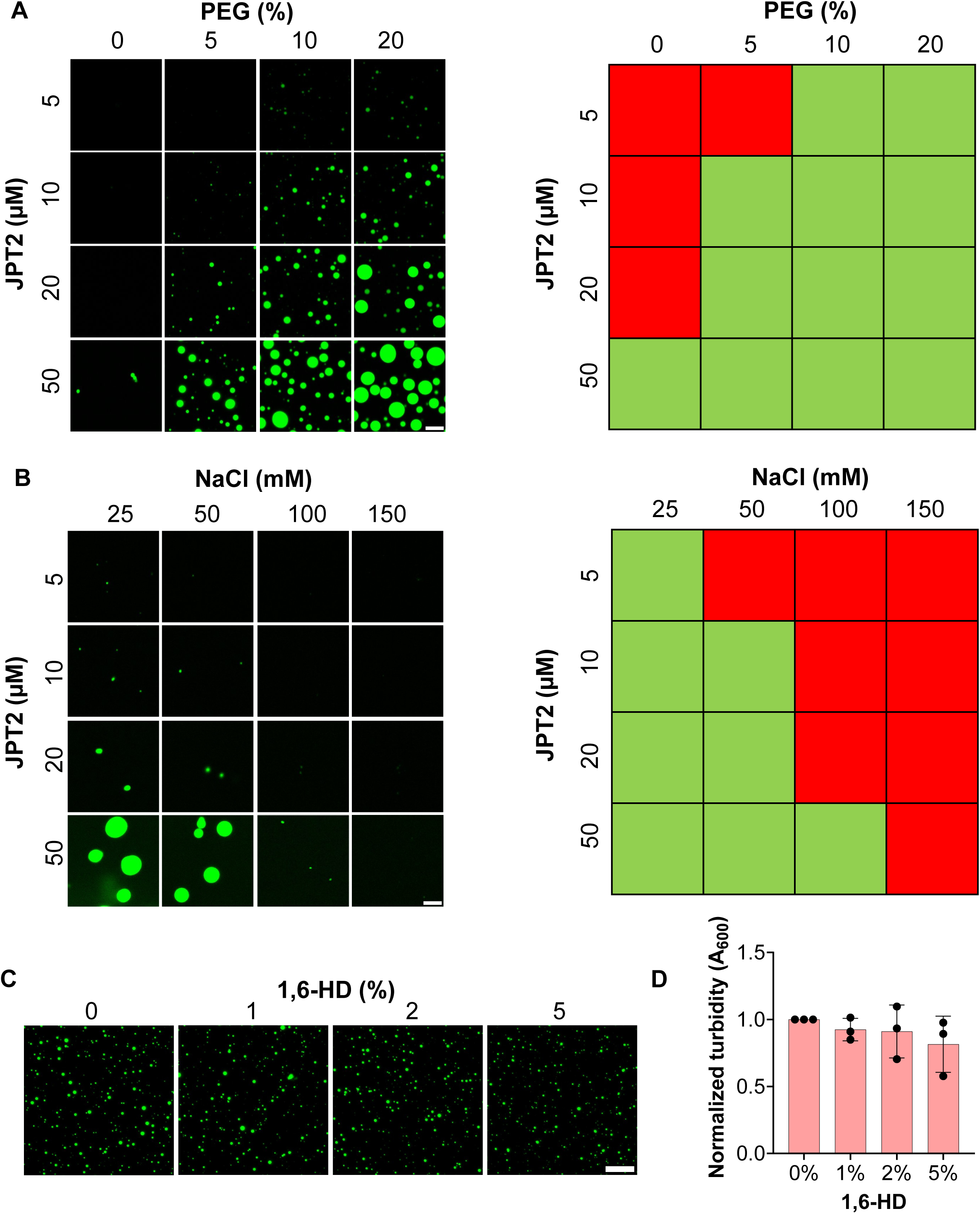
Electrostatic interactions predominantly drive JPT2 LLPS. **(A)** Representative confocal images (*left*) and phase diagram (*right*) showing JPT2 condensation with increased protein or PEG concentration. Green boxes, droplets; red boxes, no droplets. Scale bar, 5 µm. **(B)** Representative confocal images (*left*) and phase diagram (*right*) showing the effect of increasing NaCl and protein concentration on JPT2-LLPS. Green boxes, condensates; red boxes, no condensates. Scale bar, 5 µm. **(C)** Representative confocal images (*left*) showing the effect of 1,6-hexandiol (1,6-HD) on JPT2-LLPS induced by incubating 10 µM JPT2 with 10% PEG (25 mM HEPES pH 7.4, 150 mM NaCl, 10% PEG). Scale bar, 10 µm. **(D)** Turbidity assay showing absorbance (A_600_) of 10 µM JPT2 in 10% PEG (25 mM HEPES pH 7.4, 150 mM NaCl, 10% PEG) incubated with the indicated 1,6-HD concentrations for 1 hour. Data are mean ± SD from three independent experiments. In (A), (B), and (C), unlabeled JPT2 was spiked with 1% Alexa Fluor 488-conjugated JPT2, and images were captured 1 hour after LLPS-induction.

Consistent with this interpretation, addition of the aliphatic alcohol 1,6-hexanediol (≤5%), which disrupts weak hydrophobic interactions, produced only modest inhibition of JPT2 condensation as assessed by fluorescence imaging (**Fig. 4C**) and turbidity measurements (**Fig. 4D**). Collectively, these data indicate that LLPS of JPT2 is driven predominantly by electrostatic rather than hydrophobic interactions.

### JPT2 condensates display liquid-like properties

To assess whether JPT2 condensates display liquid-like behavior, droplet dynamics were examined by time-lapse confocal microscopy. JPT2 droplets exhibited surface wetting on glass coverslips, spreading upon contact and gradually losing their spherical morphology (**Fig. 5A**; **Movie S1**). Droplets in close proximity frequently fused to form larger droplets (**Fig. 5B; Movie S2**), leading to an increase in droplet size over time (**Fig. 5C**). Fluorescence recovery after photobleaching (FRAP) experiments performed on droplets formed under low-salt conditions revealed partial recovery (62.7 ± 0.2% recovery; half-time 66.9 ± 2.5 s), indicating dynamic exchange of JPT2 molecules (**Fig. 5D**). Together, these behaviors are hallmarks of liquid-like condensates and demonstrate that JPT2 undergoes LLPS *in vitro*.

**Figure 5.**
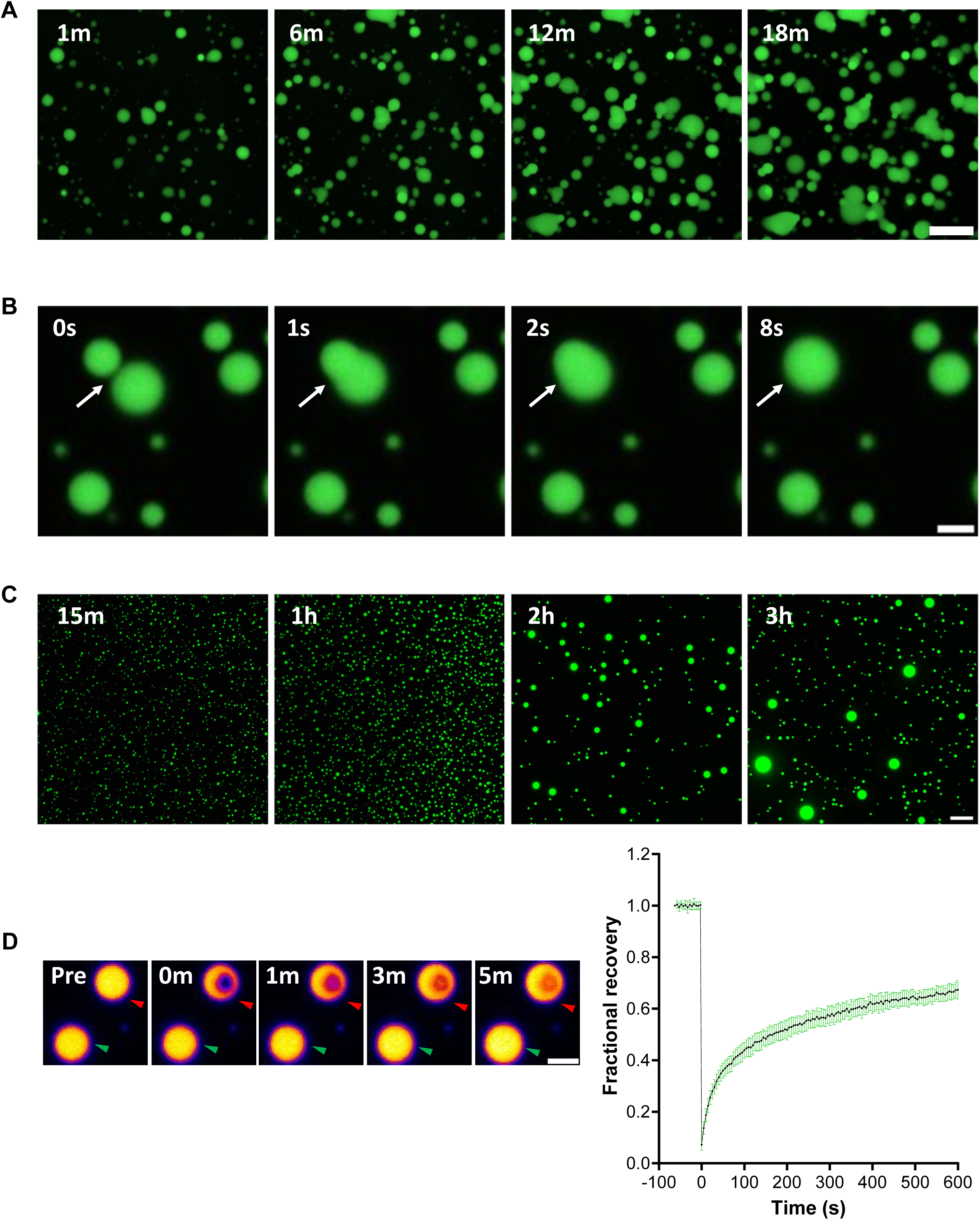
JPT2 condensates display liquid-like properties characteristic of LLPS. **(A)** Representative time-lapse confocal images showing JPT2 droplets wetting the surface of the coverglass. **(B)** Representative confocal images showing fusion of two JPT2 droplets to form a larger droplet (white arrow). **(C)** Representative time-lapse confocal microscopy images illustrating an increase in JPT2 droplet size over time. **(D)** Representative time-lapse confocal images (*left*) of a FRAP assay on JPT2 droplets. The photobleached droplet (red arrowhead) and an unbleached reference droplet (green arrowhead) are indicated. Quantification of fractional fluorescence recovery from n = 9 droplets is shown (*right*). In all assays, LLPS was induced by incubating purified JPT2 (50 µM) in a low-salt buffer (25 mM HEPES pH 7.4, 25 mM NaCl). Unlabeled JPT2 was spiked with 1% Alexa Fluor 488-conjugated JPT2. Scale bars: 10 µm (A), 2 µm (B, D), and 20 µm (C).

### LSM12 does not undergo autonomous LLPS but is recruited into JPT2 condensates

To determine whether LSM12 undergoes autonomous LLPS like JPT2, recombinant LSM12 was purified and subjected to *in vitro* LLPS assays analogous to those used for JPT2. Addition of PEG3350 (10%) to purified LSM12 resulted in the appearance of irregularly shaped structures by both DPC and fluorescent microscopy (**Fig. 6A**). Confocal imaging of fluorescently labeled LSM12 (Alexa Fluor-568) revealed that these structures increased in size, without coalescing, over time, consistent with aggregation rather than liquid-like condensation (**Fig. 6B**). Incubation of LSM12 under low-salt conditions also failed to induce condensate formation (**Fig. 6C**). Thus, unlike JPT2 (**Figs 3-5**), LSM12 does not undergo autonomous LLPS.

**Figure 6.**
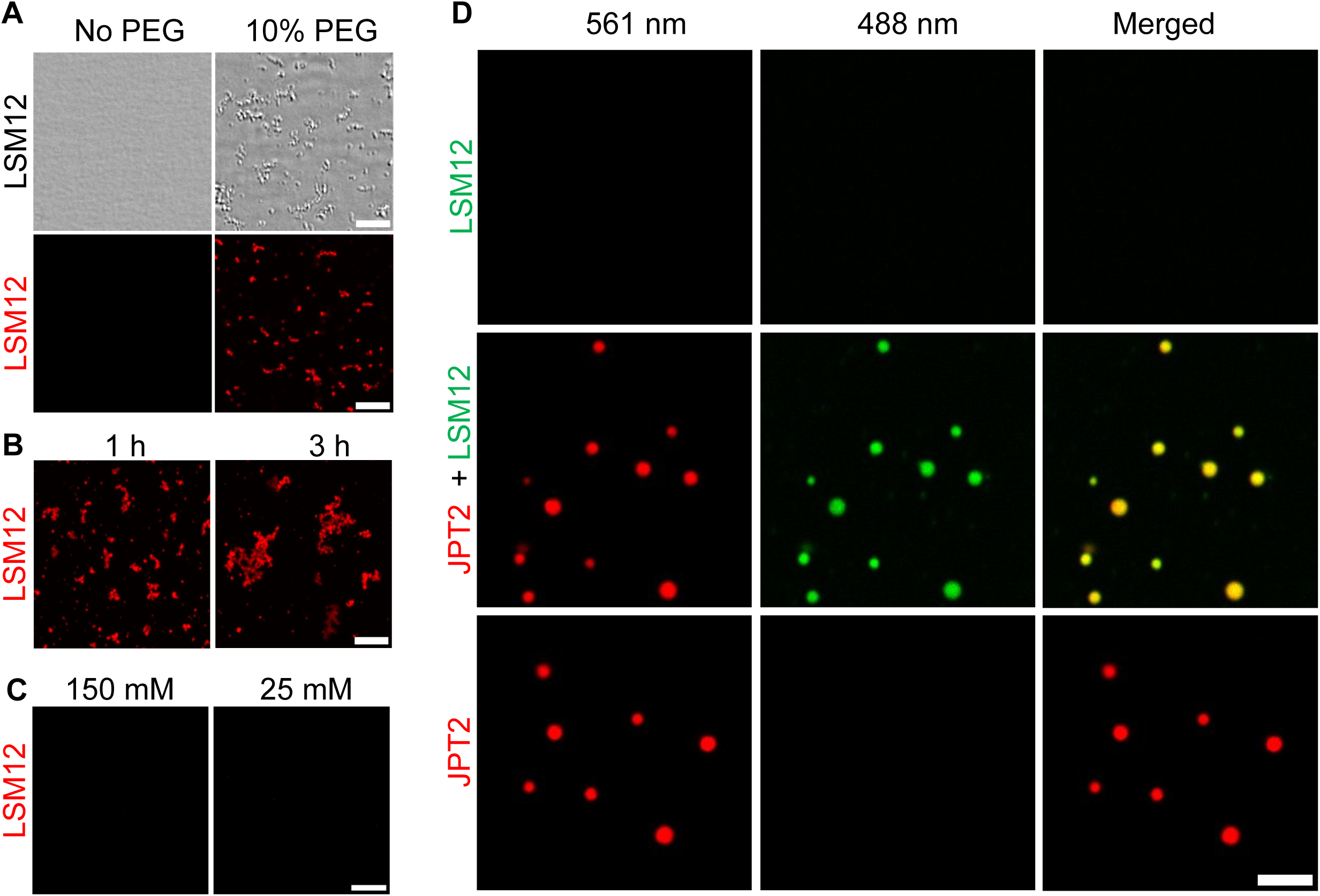
JPT2 recruits LSM12 into LLPS condensates. **(A)** Representative DPC (upper panels) and confocal microscopy images (lower panels) of purified LSM12 (20 µM) incubated for 1 hour in the presence or absence of 10% PEG3350 (25 mM HEPES, 150 mM NaCl, pH 7.4). **(B)** Representative confocal images of purified LSM12 (20 µM) incubated with 10% PEG3350 for 1 or 3 hours, illustrating time-dependent accumulation of irregular assemblies. **(C)** Representative confocal images of purified LSM12 (20 µM) incubated for 1 hour at the indicated NaCl concentrations. **(D)** Representative confocal images of LSM12 alone (25 µM*; top*), JPT2 and LSM12 co-incubated at equimolar concentrations (25 µM each; *middle*), and JPT2 alone (25 µM; *bottom*) following incubation for 1 hour in low-salt buffer (25 mM HEPES, 25 mM NaCl, pH 7.4). For (A-C), unlabeled LSM12 was spiked with 1% Alexa Fluor 568-conjugated LSM12. In (D), unlabeled JPT2 was spiked with 1% Alexa Fluor 568-conjugated JPT2 protein, and unlabeled LSM12 was spiked with 1% Alexa Fluor 488-conjugated LSM12. Scale bars: 10 µm (A), 20 µm (B, C), and 5 µm (D).

Because both JPT2 and LSM12 are required for NAADP-evoked Ca^2+^ release in mammalian cell lines [35], we next evaluated their potential interaction in the context of LLPS. Bioinformatic analyses using the PICNIC algorithm predict that LSM12 may localize to membrane-less organelles (**Fig. 1K**). In many LLPS systems, condensates are nucleated by a subset of key proteins, while additional components that do not independently phase separate can be selectively recruited to impart specific functions [60].

To test whether LSM12 can be recruited into JPT2 condensates, co-incubation assays were performed using equimolar concentrations of fluorescently labeled JPT2 (Alexa Fluor-568) and LSM12 (Alexa Fluor-488) under LLPS conditions (25 mM NaCl). Whereas LSM12 alone did not form condensates, co-incubation with JPT2 resulted in recruitment of LSM12 into JPT2 droplets, as evidenced by strong co-localization (partition coefficient ∼4, **Fig. 6D**). These data indicate that although LSM12 does not autonomously undergo LLPS, it can partition into JPT2 condensates.

### JPT2 condensates sequester NAADP

To examine NAADP dynamics in the context of JPT2 phase separation, two fluorescent analogs of NAADP were synthesized (**Fig. S3**): 5-(AZ488-[CH_2_]_6_)-NAADP (AZ488-NAADP) and BODIPY-FL-(EG_4_)-NAADP (BODIPY-NAADP). To test activity, binding of these analogs to JPT2 was first assessed using biolayer interferometry (BLI) [35]. AZ488-NAADP associated with JPT2 (**Fig. 7A & S4A**), whereas BODIPY-NAADP showed no detectable interaction (**Fig. 7B**). Importantly, unconjugated AZ488 did not bind JPT2 (**Fig. S4B**), indicating that the interaction observed with AZ488-NAADP was not driven by the fluorophore itself.

**Figure 7.**
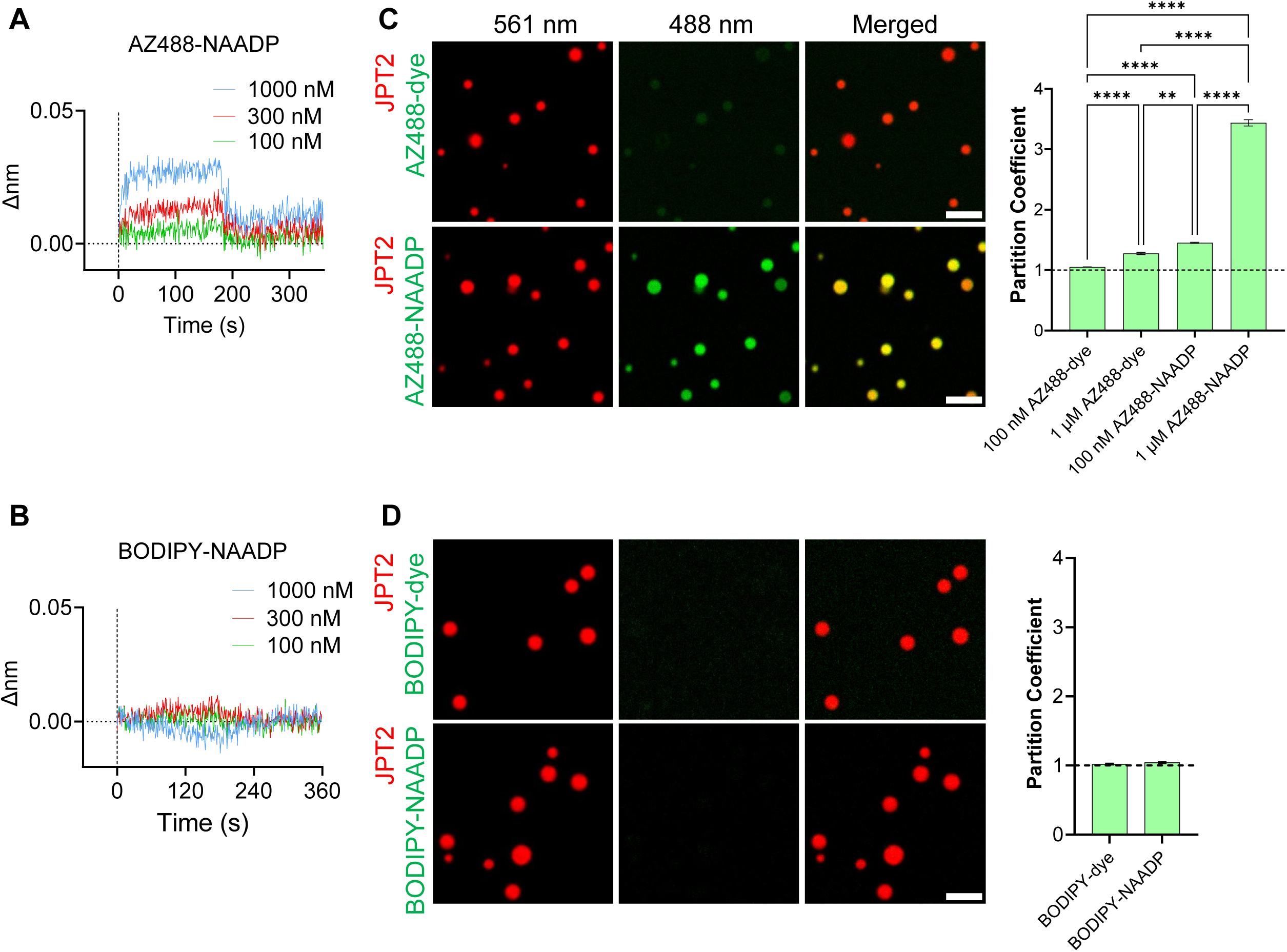
JPT2 condensates selectively bind and sequester AZ488-NAADP *in vitro*. **(A**) Representative BLI traces showing association and disassociation kinetics of AZ488-NAADP binding to recombinant JPT2. **(B)** Representative BLI traces showing a lack of detectable interaction between BODIPY-NAADP and recombinant JPT2. **(C)** Representative confocal images (*left*) of JPT2 condensates co-incubated with AZ488-dye (1 µM, *upper panels*) or AZ488-NAADP (1 µM, *lower panels*) in low salt buffer (25 mM HEPES, 25 mM NaCl, pH 7.4). Bar graph (*right*) quantifying enrichment of the indicated concentrations of AZ488-dye or AZ488-NAADP within JPT2 condensates. Data are shown as mean ± SD; n ≥ 30 droplets, ****p<0.0001, **p<0.01 (unpaired t-test). **(D)** Representative confocal images (*left*) of JPT2 condensates co-incubated with BODIPY-dye (100 nM, *upper panels*) or BODIPY-NAADP (100 nM, *lower panels*) under low-salt conditions (25 mM HEPES, 25 mM NaCl, pH 7.4). Bar graph (*right*) quantifying enrichment of BODIPY-dye or BODIPY-NAADP within JPT2 condensates. Data shown are mean ± SD; n=20 droplets. Scale bars, 5 µm.

These fluorescent NAADP analogs were next examined in phase separation assays. AZ488-NAADP was enriched within JPT2 condensates, as evidenced by increased fluorescence within droplets relative to the unconjugated fluorophore control (**Fig. 7C**). In contrast, BODIPY-NAADP, did not partition into JPT2 condensates, with conjugated and unconjugated fluorophores displaying comparable partition coefficients (**Fig. 7D**).

To assess whether NAADP partitioning reflected specific interactions with JPT2 rather than nonspecific accumulation within condensates, recruitment of AZ488-NAADP into FUS-condensates was examined. Recombinant MBP-FUS was induced to phase separate by TEV protease treatment, followed by the addition of AZ488-NAADP or unconjugated AZ488 (100 nM). AZ488-NAADP did not show preferential recruitment into FUS condensates relative to unconjugated AZ488 (**Fig. S4C**), in contrast to the selective partitioning of AZ488-NAADP observed in JPT2 droplets. Finally, to assess the reversibility of AZ488-NADDP sequestration into droplets, assays were performed with unlabeled NAADP. Addition of excess unlabeled NAADP inhibited AZ488-NAADP labeling of JPT2 condensates (**Fig. S5**), showing compartmentalized NAADP was exchangeable with NAADP added to the media.

Collectively, these results demonstrate that a fluorescent NAADP analog capable of binding JPT2 selectively partitions into JPT2 condensates *in vitro*, whereas an analog that does not bind JPT2 does not. NAADP sequestration was specific to JPT2 condensates and is not a general feature of phase-separated assemblies.

### JPT2 condensates associate with microtubules

Because JPT2 is a known microtubule binding protein [61–63], we examined the behavior of JPT2 condensates in the presence of polymerized microtubules. Microtubules were assembled and stabilized with taxol in polymerization buffer (see ‘Methods’), then diluted into low- or high-salt buffers in the absence or presence of JPT2. As observed previously, JPT2 condensates formed under low salt but not high salt conditions (**Fig. 8 A,B**). In the absence of JPT2, microtubules were detectable under both salt conditions, although their number and length were reduced at higher NaCl concentrations (**Fig. 8 C,D**), consistent with the known inhibitory effects of elevated ionic strength on microtubule assembly [64].

**Figure 8.**
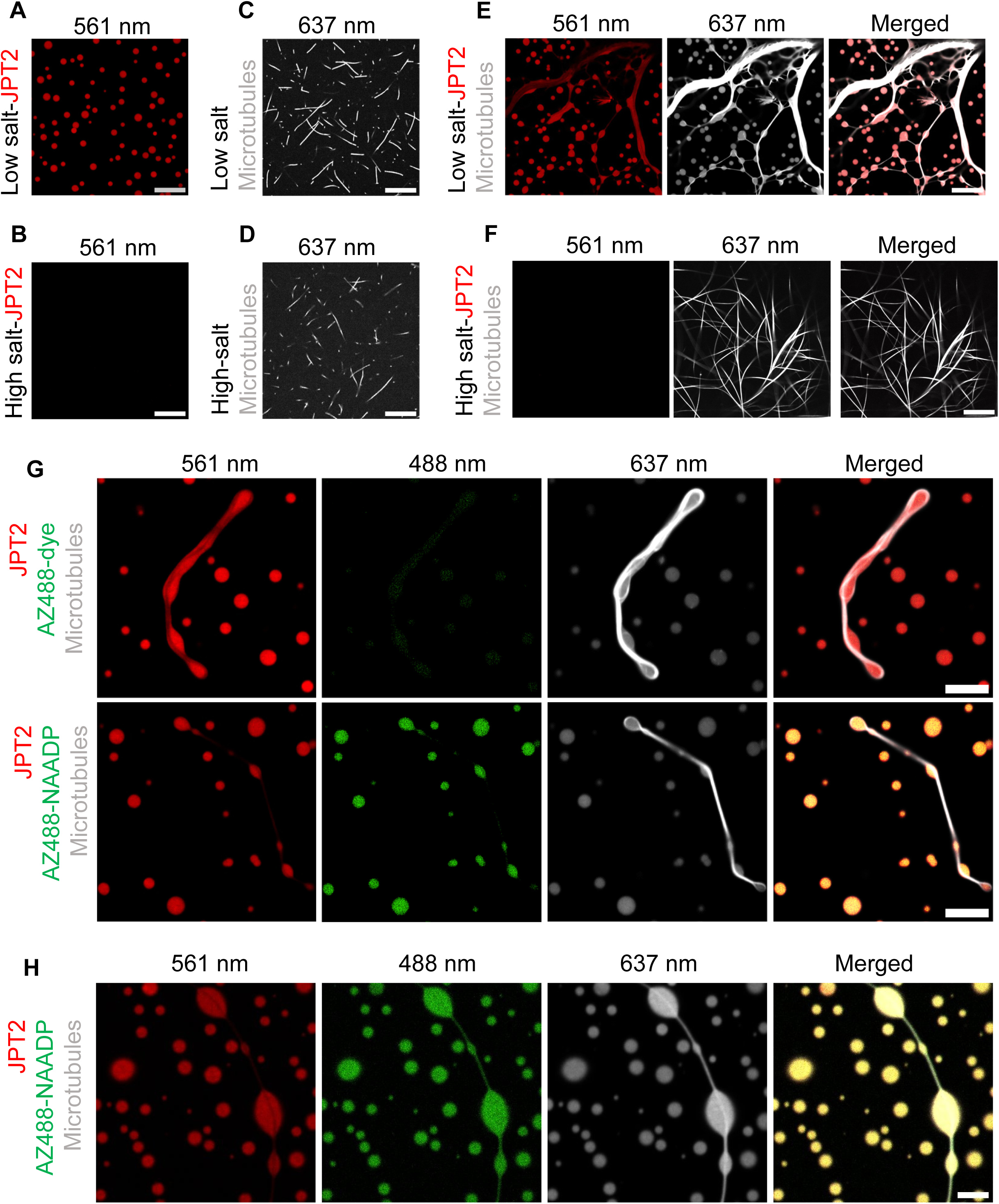
JPT2 condensates associate with microtubules and lysosomes *in vitro*. **(A, B)** Representative confocal images of purified JPT2 protein incubated in low salt buffer (A) or high salt buffer (B). **(C, D)** Representative confocal images of polymerized microtubules incubated in low salt buffer (C) or high salt buffer (D). **(E)** Representative confocal images of pre-formed JPT2 condensates co-incubated with microtubules in low salt conditions. **(F)** Representative confocal images of purified JPT2 co-incubated with microtubules in high-salt conditions. **(G)** Representative confocal images of pre-formed JPT2 condensates co-incubated microtubles in low-salt buffer in the presence of AZ488-dye or AZ488-NAADP (100 nM). **(H)** Magnified images to show AX488-NAADP tracking JPT2 distribution along the length of a microtubule. Images were captured in low salt buffer, 30 mins after mixing components.

We next examined samples in which preformed JPT2 condensates and microtubules were co-incubated. Under low salt conditions, JPT2 condensates readily formed and fluorescently labeled tubulin was enriched within these condensates, as evidenced by fluorescence colocalization (**Fig. 8E**). Individual JPT2 condensates frequently associated with microtubule structures, often decorating a single microtubule, in a ‘beads-on-a-string’-like arrangement (**Fig. 8E**). Condensates were commonly observed at microtubule branch points or regions of curvature (**Fig. 8E**). Whereas free JPT2 droplets were spherical, condensates associated with the microtubules lost spherical morphology and spread into irregular shapes along the microtubule surface (**Fig. 8E**), reminiscent of the surface-wetting behavior observed on uncoated coverglass (**Fig. 4A**).

Finally, these assays were repeated in the presence of AZ488-NAADP. Addition of AZ488-NAADP did not affect the association of JPT2 condensates with microtubules (**Fig. 8G**). Instead, AZ488-NAADP partitioned into JPT2 condensates and was delivered into close proximity with microtubules, mirroring the spatial distribution of JPT2. The unconjugated fluorophore AZ488 did not show this behavior (**Fig. 8G**). In high resolution images, the distribution of AZ488-NAADP tracked along with JPT2 throughout the length of microtubule (**Fig. 8H**). Together, these data indicate that JPT2 condensates, and their cargo, associate with microtubules through the intrinsic microtubule-binding properties of JPT2 [61–63].

### JPT2 condensates associate with lysosomes

Finally, we examined the relationship between JPT2 condensates and lysosomes. Lysosomes isolated from HEK293 cells and labeled with LysoTracker Deep Red were mixed with pre-formed JPT2 condensates formed under low salt conditions. Confocal microscopy performed 15-30 minutes after mixing revealed frequent association of lysosomes with JPT2 condensates (**Fig. 9A**). This interaction was not characterized by complete engulfment or simple peripheral association, but rather as a partial overlap at the perimeter of the condensate (**Fig. 9A, inset**). Quantification analysis revealed that approximately 40% of lysosomes were associated with JPT2 condensates under these incubation conditions (**Fig. 9B**), and individual JPT2 droplets were often observed to associate with multiple lysosomes. These observations demonstrate that JPT2 condensates can also interact with lysosomes *in vitro*.

**Figure 9.**
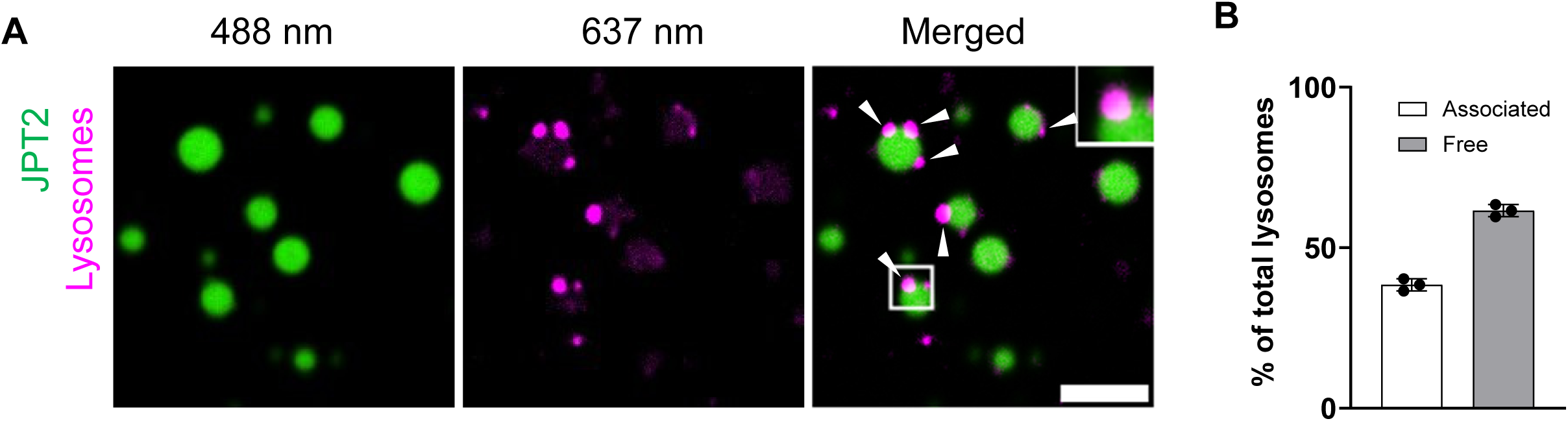
Interaction between JPT2 condensates and lysosomes. **(A)** Representative confocal images showing association of isolated lysosomes with JPT2 condensates (white arrowheads). Inset shows partial overlap at the phase boundary between a JPT2 condensate and an associated lysosome. **(B)** Quantification of the fraction of free verses JPT2 condensate-associated lysosomes. Data are shown as mean ± SD; n=3. Purified JPT2 protein was labeled with Alexa Fluor 568 in (A, B, E-G), and with Alexa Fluor 488 in (H). Microtubules in (C-G) were assembled from Hilyte-647-labeled tubulin. Lysosomes in (H) were isolated from HEK293 cells and labeled with Lysotracker Deep Red. Scale bars: 20 µm (A-F). 10 µm (G), 5 µm (H).

## Discussion

Collectively, these findings demonstrate that the intrinsically disordered protein JPT2 undergoes liquid-liquid phase separation under conditions of molecular crowding and/or reduced ionic strength *in vitro*. The resulting JPT2 condensates selectively sequester NAADP, recruit the NAADP-binding protein LSM12, associate with lysosomes, and interact with microtubules, influencing their organization under phase separating conditions. Together, these properties suggest that phase separation of JPT2 provides a plausible organizational framework for coordinating multiple components of the NAADP signaling pathway, and advance two important avenues for understanding NAADP action.

First, our data raise fundamental questions about how NAADP binds with high affinity to an entirely disordered protein lacking any defined tertiary structure. Yet high affinity interactions involving intrinsically disordered regions (IDRs) within proteins, or IDPs, are increasingly well documented, with reported affinities spanning picomolar to micromolar ranges [65–70]. Disorder is prevalent within the human proteome, and confers distinctive regulatory features, such as enhanced binding sensitivity, conformational flexibility, multivalency, and promiscuity, compared with classical ‘lock-and-key’ interactions characteristic of structured proteins [71, 65, 72]. In the context of JPT2, it will be important to define how NAADP binding is achieved at the molecular level and how this relates to interactions with different ion channel targets including TPCs [31, 35], ryanodine receptors [32], as well as other cellular effectors.

Second, the propensity of JPT2 to undergo LLPS introduces a potentially new regulatory dimension for choreographing NAADP-evoked Ca^2+^ signaling. Whereas Ca^2+^ signaling has traditionally been conceptualized in four dimensions (space and time, xyz-t), phase separation introduces an additional level of complexity by partitioning signaling components into a demixed phase distinct from the bulk cytoplasm. Condensates possess physical and chemical properties that are different from the dilute phase, enabling selective concentration of ions, small molecules, and proteins, as well as altered diffusion and buffering characteristics [73, 74]. Notably, recent work in bacteria has demonstrated that condensates can sustain intracellular Ca^2+^ concentrations ∼5-fold higher than the surrounding cytoplasm, with consequences for membrane potential, gene expression, and ion homeostasis [75]. In vertebrate cells, biomolecular condensates play a critical role in cytoplasmic organization with ∼18% of the vertebrate proteome organized as condensates [76], driving an increased recognition of their significance for cell physiology and disease [74].

Although our data derive from *in vitro* analyses, the observation that both JPT2 and LSM12 can localize to condensate-like assemblies motivates future investigation of NAADP-BP phase behavior in cells. This idea is supported by systems-level datasets reporting enrichment of these proteins in membrane-less organelles [77–79], as well as a growing literature implicating phase separation in multiple aspects of NAADP biology. These include enzymatic generation of nucleotide-based second messengers [80], organization of membrane contact sites [81–83], the clustering and local regulation of ion channels [84, 85] as well as control of endolysosomal signaling processes linked to NAADP [86–92].

How might phase separation influence NAADP-evoked Ca^2+^ release? Several long-recognized but poorly understood features of NAADP signaling could be explained by LLPS of NAADP-BPs. Compartmentalization of JPT2 and LSM12 into condensates could potentially serve inhibitory or facilitatory roles depending on context. For example, sequestration of NAADP-BPs away from TPCs would be expected to reduce cellular responsiveness to NAADP, providing a potential ‘off’ mechanism. This may contribute to cell-type-specific variability in NAADP sensitivity and highlights the importance of examining NAADP-BP distribution and dynamics across different cellular states. Phase separation may also operate on shorter timescales to attenuate NAADP signaling. Early studies in sea urchin egg homogenate revealed a striking self-inactivation phenomenon, whereby irreversible binding of ^32^P-NAADP led to desensitization even at low ligand concentrations. If NAADP binding promotes condensation of NAADP-BPs, and the concentration of NAADP-BPs in condensates exceeds local NAADP concentrations, this provides an obvious mechanism to prevent subsequent addition of NAADP from activating TPCs. Reversal of desensitization would then require condensate dissolution, providing a mechanism by which the kinetics and competence of NAADP signaling could by dynamically regulated. Transitioning NAADP-BPs into and out of phase could define the kinetics of NAADP action, and even whether cells are competent to respond to NAADP at all.

Conversely, our data also support a facilitatory role for JPT2 phase separation. JPT2 condensates selectively concentrate NAADP (**Fig. 7**), recruit LSM12 (**Fig. 6**), associate with lysosomes (**Fig. 9**), and interact with microtubules (**Fig. 8**), thereby co-localizing multiple components essential for NAADP-evoked Ca^2+^ release. Autonomous LLPS by JPT2 likely reflects intrinsic sequence features characteristic of phase separating proteins, including extensive disorder, low hydropathy, high net charge (predicted pI ∼9.4), and multivalency (**Fig. 1**). Consistent with this, JPT2 phase separation is strongly sensitive to ionic strength, indicating a dominant role for electrostatic interactions (**Figs. 3 & 4**). This suggests a potential regulatory mechanisms *in vivo*, such as charge-altering post-translational modifications, localized pH changes, or ion fluxes generated by nearby ion channels. In contrast, the limited effect of 1,6-HD indicates a comparatively minor contribution for hydrophobic interactions to JPT2 condensation *in vitro* (**Fig. 4**). The physicochemical environment within JPT2 condensates supports recruitment of client molecules, including LSM12 (**Fig. 6**) and NAADP (**Fig. 7**), consistent with known principles of condensate biology where scaffold proteins nucleate assemblies that subsequently incorporate additional components to modulate function and localization. In this way, JPT2 condensates may serve as microdomains that integrate multiple activatory cues required for NAADP signaling.

Phase separation of NAADP-BPs may also facilitate spatial targeting of NAADP signaling to membrane contact sites between acidic Ca^2+^ stores and the ER, where NAADP-dependent crosstalk between TPCs and ER Ca^2+^ release channels occurs [93]. Because cytoplasmic diffusion of NAADP would be expected to activate TPCs broadly, mechanisms that restrict signaling to specialized subcellular zones are likely required. JPT2 may contribute to such targeting through its ability to bind both endolysosomal channels and the microtubule network that parallels ER organization. Phase separation, potentially promoted by molecular crowding at these contact sites where diffusion is constrained [83], could further enable localized membrane wetting [94] and microtubule decoration by JPT2 (**Fig. 8**), thereby defining spatially restricted NAADP signaling platforms [95, 81, 96, 89–92].

Finally, the microtubule binding capacity of JPT2 likely plays a key role. When JPT2 condensates associate with microtubules, they lose spherical droplet morphology and spread along the filament, decorating the structures with a NAADP-BP (**Fig. 8**). The microtubule-binding property of JPT2 was originally reported in *Drosophila* [61], but is also apparent in human cells (Fig. 8, [62, 63]) and is mediated by the repeat PPGGK*S motifs within the JPT family [62]. This behavior parallels membrane wetting and resembles the properties of other microtubule-associated proteins, including Tau. Like JPT2, Tau undergoes electrostatically phase separation [52], and Tau-tubulin condensates have been proposed to organize microtubules through multiple mechanisms [97, 98]. The effects of JPT2 on microtubule morphology observed here (**Fig. 8**), together with prior cellular studies [61–63], support the classification of JPT2 as a *bona fide* microtubule associated protein and suggest additional roles in microtubule organization and dynamics that warrant further investigation.

In summary, our data identify phase separation of NAADP binding proteins as a previously unrecognized mechanism with the potential to organize NAADP signaling components in space and time. These findings motivate future studies to define when and where NAADP-BP condensates form in cells and how their physical properties contribute to regulation of Ca^2+^ signaling under physiological and pathological conditions.

## Data availability

All data and protocols within this manuscript will be shared upon reasonable request.

## Author contributions

All authors were involved in conceptualization and design of the research. Performed research and analyzed data: SK, GSG, JC, KS, MM, ZG. Wrote the manuscript: SK & JSM, with input from all authors. All authors reviewed the final version of the manuscript.

## Acknowledgements

This work was supported by NIH grant number GM088790 and S10 OD025000JC. We appreciate the use of the Oxford Instruments Center for Advanced Microscopy - Electron Microscopy Core (RRID:SCR_026315) at MCW for assisting with imaging experiments, as well as the MCW Program in Chemical Biology. JC was supported by a Steel Perlot Early Investigator Grant and Erika Alden DeBenedictis. KS was supported by the University College London-Birkbeck Medical Research Council Doctoral Training Programme (MR/W006774/1). GTH was supported by a BBSRC Discovery Fellowship (BB/X009955/1). Access to ultra-high field NMR spectrometers was supported by the Francis Crick Institute through provision of access to the MRC Biomedical NMR Centre. The Francis Crick Institute receives its core funding from Cancer Research UK (FC001029), the UK Medical Research Council (FC001029), and the Wellcome Trust (FC001029). The graphical abstract was created using BioRender.

