## Supplementary Figures for "Liquid liquid phase separation of the intrinsically disordered protein JPT2 compartmentalizes components of NAADP-evoked Ca^2+^signaling"

**Supplementary Material**

**Figure S1.** Turbidity assay showing absorbance ($A_{600}$) of purified JPT2 (20 µM) incubated for 1 hour at the indicated NaCl concentrations. Data are shown as mean ± SD from three independent experiments. Statistical significance was assessed by one-way ANOVA (****p<0.0001, ***p<0.001, *p<0.05).

**Figure S2.** Box-and-whisker plot showing the distribution of JPT2 droplet size formed by incubating purified JPT2 (50 µM) in low salt buffer (25 mM HEPES, 25 mM NaCl, pH 7.4) or under molecular crowding conditions (25 mM HEPES, 150 mM NaCl, 10% PEG3350, pH 7.4). Boxes indicate median and interquartile range; whiskers denote minimum and maximum values. n = 220 droplets per condition.

**Figure S3.** Chemical structures of **(A)** AZ488-NAADP and **(B)** BODIPY-NAADP.

**Figure S4.** Representative BLI traces showing the association and dissociation of **(A)** AZ488-NAADP or **(B)** AZ488-dye with recombinant JPT2. **(C)** Representative confocal microscopy images (*left*) showing phase separated FUS droplets co-incubated with AZ488-dye (upper panels) or AZ488-NAADP (lower panels). Scale bar, 5 µm. Bar graph (*right*) quantifying enrichment of AZ488-dye and AZ488-NAADP in FUS condensates in low-salt conditions. Data are shown as mean ± SD; n=10 droplets.

**Figure S5.** Representative confocal images (*left*) of JPT2 condensates co-incubated with 1 µM AZ488-dye (*upper panels*), 1 µM AZ488-NAADP (*middle panels*), or 1 µM AZ488-NAADP with 1 mM unlabeled NAADP (lower *panels*) in low salt buffer (25 mM HEPES, 25 mM NaCl, pH 7.4). Bar graph (*right*) quantifying enrichment of AZ488-dye or AZ488-NAADP within JPT2 condensates. Data are shown as mean ± SD; n ≥ 24 droplets, ****p<0.0001 (one-way ANOVA).

**Movie S1.** Time-lapse movie (18 frames per second) showing JPT2 droplets, formed under low-salt conditions (25 mM HEPES, 25 mM NaCl, pH 7.4), wetting the surface of uncoated coverglass and gradually losing their spherical morphology over time.

**Movie S2.** Time-lapse movie (10 frames per second) showing fusion of JPT2 droplets formed under low-salt conditions (25 mM HEPES, 25 mM NaCl, pH 7.4).

#### Figure S1

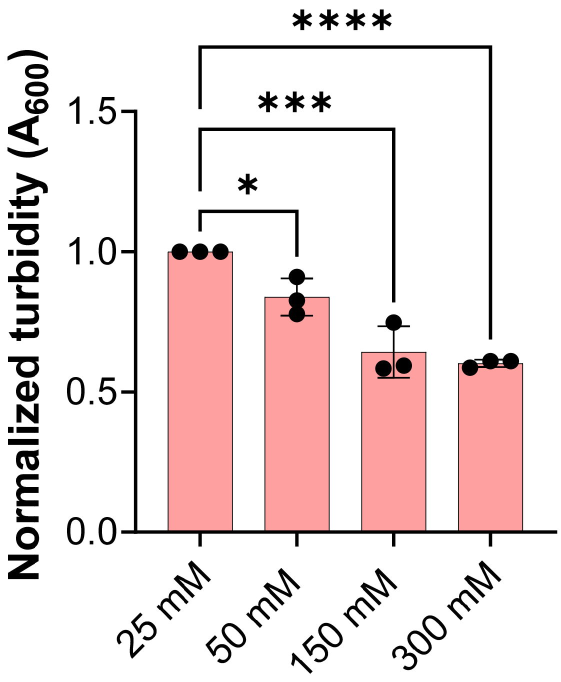

**Figure S2**

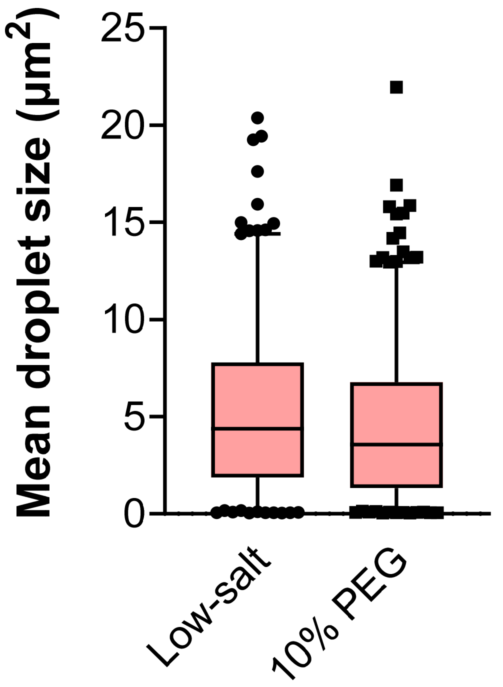

### Figure S3

**A**

5-(AZ488-$[CH_2]_6$)-NAADP

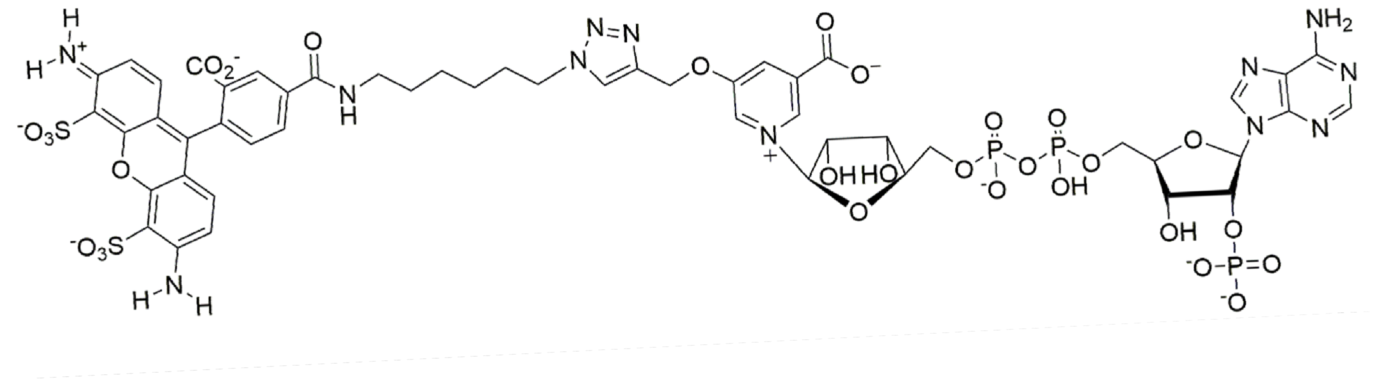


**B**

BODIPY-FL-($EG_4$)-NAADP

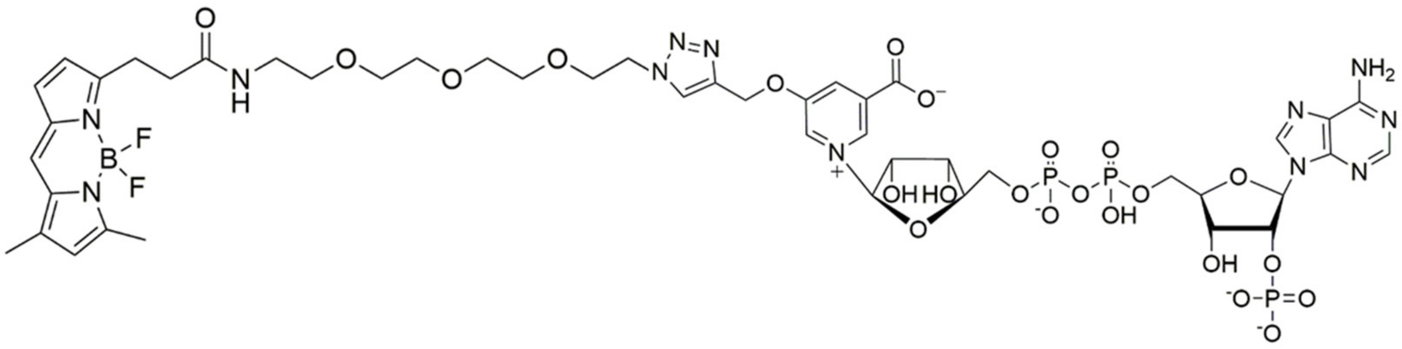

### Figure S4

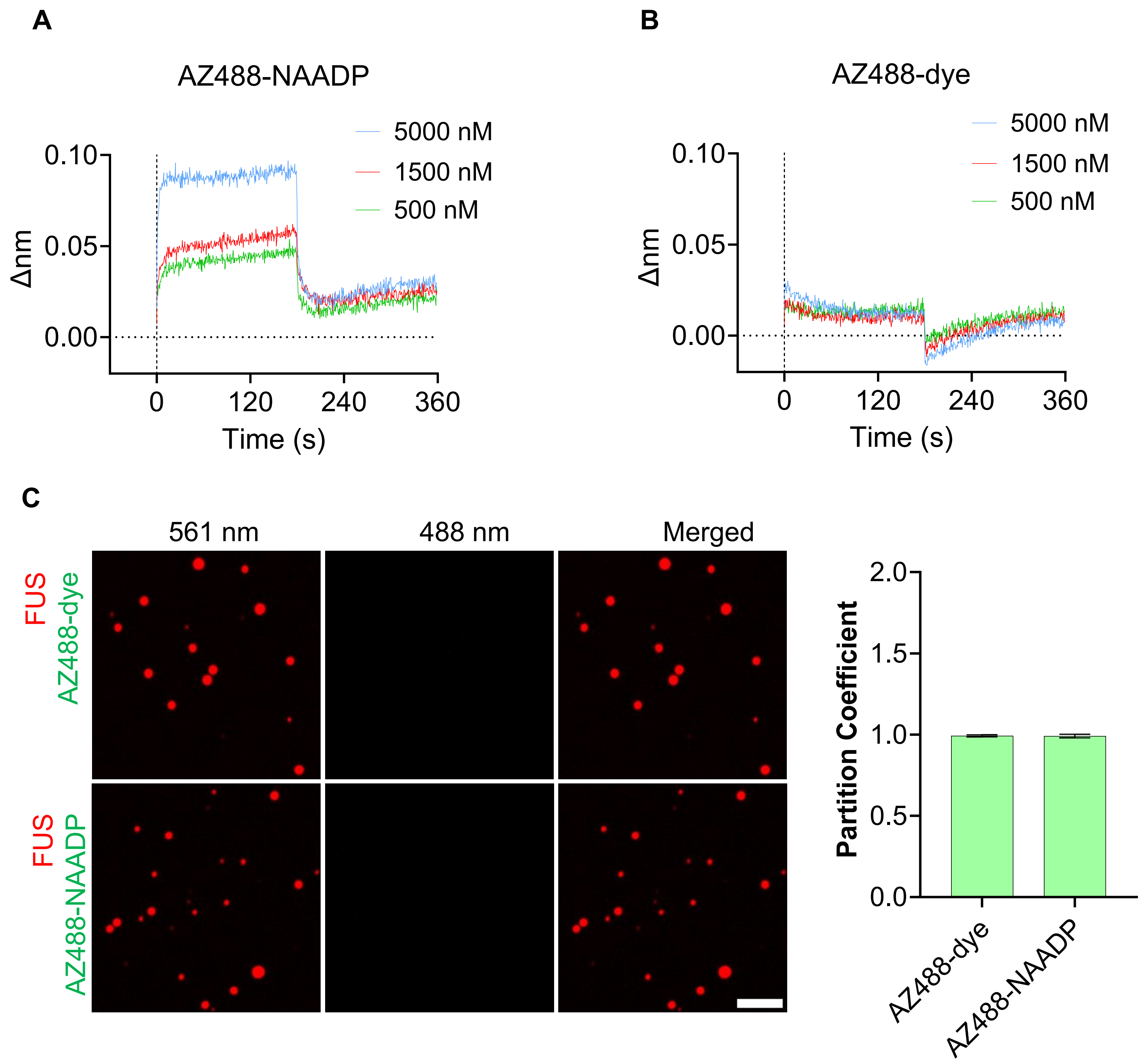

A
AZ488-NAADP
5000 nM
1500 nM
500 nM
Δnm
0.10
0.05
0.00
0
120
240
360
Time (s)
B
AZ488-dye
5000 nM
1500 nM
500 nM
Δnm
0.10
0.05
0.00
0
120
240
360
Time (s)
C
561 nm
488 nm
Merged
FUS
AZ488-dye
FUS
AZ488-NAADP
Partition Coefficient
2.0
1.5
1.0
0.5
0.0
AZ488-dye
AZ488-NAADP

#### Figure S5

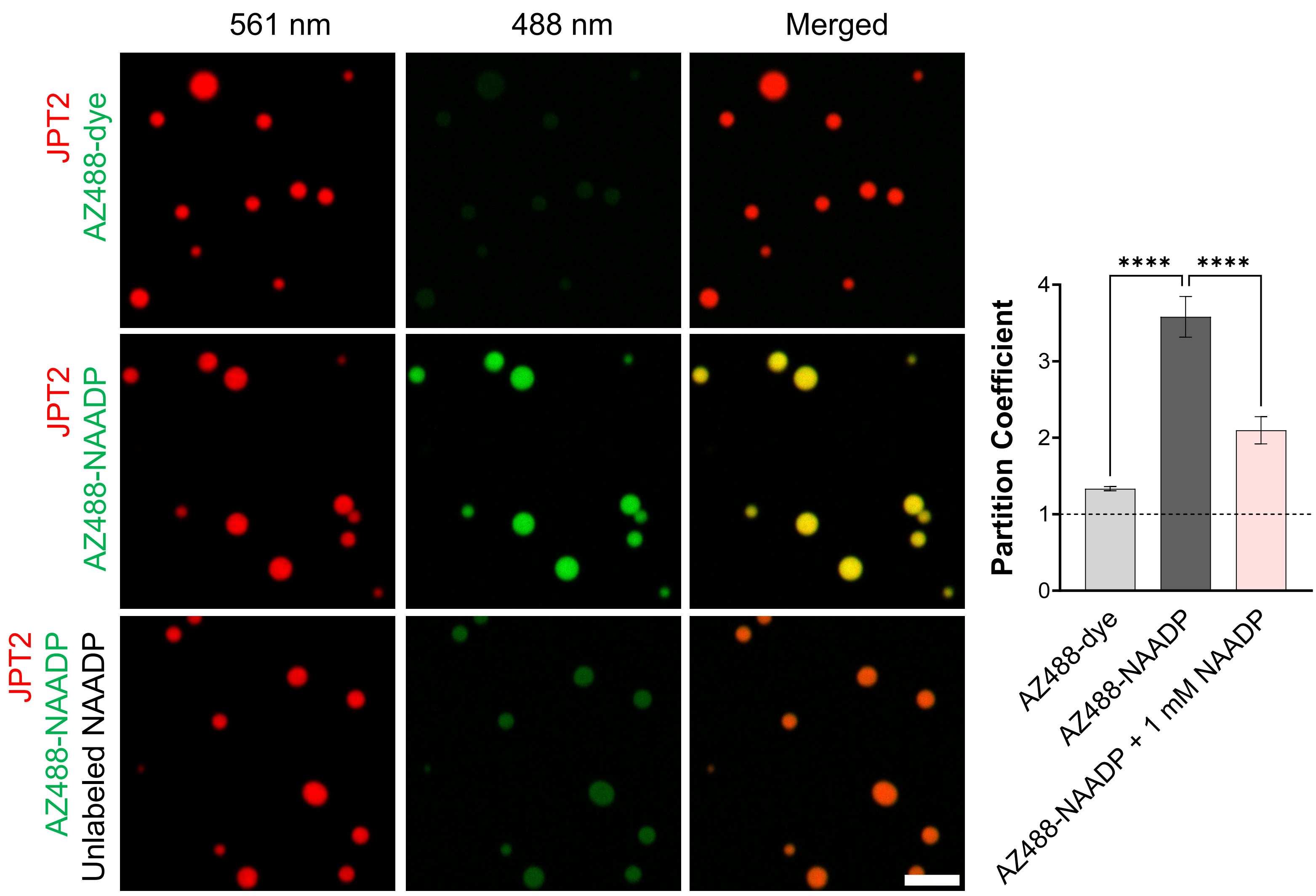
561 nm
488 nm
Merged
JPT2
AZ488-dye
JPT2
AZ488-NAADP
JPT2
AZ488-NAADP
Unlabeled NAADP
****
****
Partition Coefficient
4
3
2
1
0
AZ488-dye
AZ488-NAADP
AZ488-NAADP + 1 mM NAADP
