## Supplementary Methods for "Liquid liquid phase separation of the intrinsically disordered protein JPT2 compartmentalizes components of NAADP-evoked Ca^2+^signaling"

**Chemoenzymatic synthesis of** **long wavelength fluorescent NAADP derivatives BODIPY-NAADP and** **AZ488-NAADP.**

Detailed synthetic procedures for the production and characterizations of all new compounds and key intermediates for the synthesis of long wavelength fluorescent NAADP derivatives (**A**), AZ488-NAADP and (**B**) BODIPY-NAADP are given below.

We describe linking the acetylenic group of the NAADP analog, 5-(prop-2-yn-1-yloxy)-NAADP (**4**), with an azide group covalently connected to a BODIPY or to a AZDye488^®^ using a copper catalyzed click reaction. Linking the dye to the dinucleotide using the click reaction is high yielding, highly specific for both the alkyne and azido groups, and is compatible with aqueous-organic solvent mixtures necessary to accommodate the solubilities of both the pyridine dinucleotide and a hydrophobic dye [1]. The click reactions resulting in the synthesis of the dye conjugates are illustrated in Schemes S4 and S5. The clickable NAADP derivative **4** was synthesized using the enzyme catalyzed pyridine base exchange reaction between NADP and the nicotinic acid analog 5-(prop-2-yn-1-yloxy)-nicotinic acid (**3**) as illustrated in Scheme S2. The *Aplysia californica* ADP-ribosyl cyclase [2] catalyzed base exchange reaction was previously utilized for the synthesis of NAADP and pyridine base substituted NAADP analogs [3, 4]. Although nicotinic acid derivative **3** required for the synthesis of NAADP analog **4** was reported in a dissertation [5], it is neither commercially available nor has it been described in the peer reviewed chemical literature previously. Nicotinic acid analog **3** was synthesized in three steps from 5-hydroxynicotinic acid as illustrated in Scheme S1.

**Preparative anion-exchange chromatography**. Anion exchange chromatography was used for in the purification of NAADP analog **4**, BODIPY-NAADP (**5**) and AZ488-NAADP (**6**). Procedures for automated anion exchange separations using Bio-Rad AG MP-1 resin (BioRad Laboratories, Hercules, CA, USA) as the stationary phase in water-trifluoroacetic acid gradients was previously described [6, 7]. Open column chromatography used DEAE cellulose (DEAE Sephacel, 17050001, Cytiva, Uppsala, Sweden) packed into a 1.5 × column and equilibrated with 10 mM NH_4_HCO_3_ at pH 7.5. Samples of dinucleotides were adjusted to pH 7.5, diluted to low ionic strength, applied to the column, and washed with one column volume of 0.01 mM NH_4_HCO_3_ buffer. Once the sample had been absorbed the separation was developed by applying a linear gradient formed between 600 mL of 0.01 mM NH_4_HCO_3_ and 600 mL of 400 mM NH_4_HCO_3_. Fractions (10 mL) were collected and the absorbance of the dinucleotide determined at wavelengths corresponding to their absorption maxima. Fractions containing the product dinucleotides were combined and repeatedly lyophilized to remove the volatile NH_4_HCO_3_ buffer.

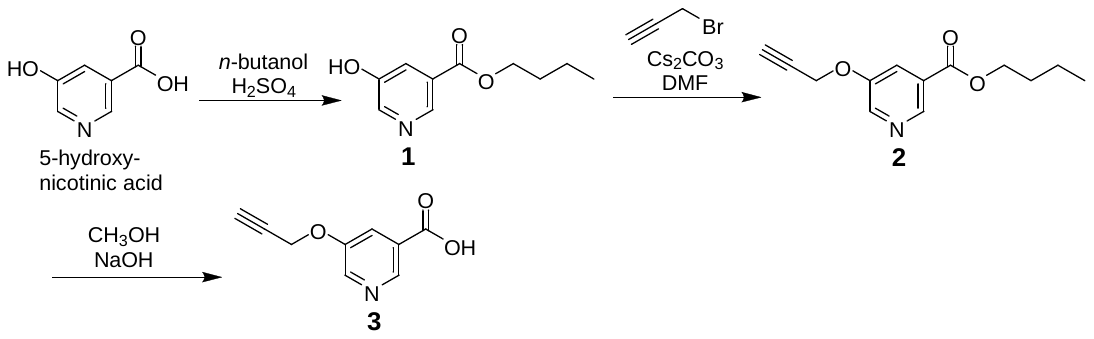

Scheme S1. Synthesis of 5-(prop-2-yn-1-yloxy)nicotinic acid (**3**) starting from 5-hydroxynicotinic acid

**Butyl 5-hydroxynicotinate** (**1**) 5-Hydroxynicotinic acid (5 g, 36.14 mmol) was dissolved in 100 mL of *n*-butanol. Concentrated H_2_SO_4_ (4 mL) was added in parts while constantly stirring. The resulting solution was heated to reflux for 48 hours with constant stirring. The reaction solution was cooled to room temperature and concentrated by distillation of the solvent *in vacuo*. NaHCO_3_ (25.2 g, 0.3 mol) and ethyl acetate (200 mL) were added to the concentrated residue and stirred vigorously for 2 hours. After 2 hours, the color of the reaction mixture changed from bright yellow to off-white and the mixture was transferred to a separatory funnel. The phases were mixed thoroughly and organic layer was washed with 3 times with100 mL portions of water and dried over anhydrous Na_2_SO_4_. Removal of the solvent by distillation *in vacuo* afforded an off-white powder which was further purified by crystallizing from CH_2_Cl_2_, affording white round crystals. (4.43 g, 63.1%). m.p. 108-110 °C; (reported m.p. 109-111 °C [5]). TLC R*_f_* 0.56 (1:1 hexane-ethyl acetate).

An analytical sample was prepared by bulb-to-bulb distillation of a 2 g sample (0.314 Torr/195-200 °C) followed by dissolving a small portion (100 mg) in boiling methanol (0.5 mL), and adding hot water (ca. 1 mL) followed by rapid cooling. White crystals (75 mg) were obtained: m.p. 125-126 °C.

Anal. Calcd. for C_10_H_13_NO_3_: C, 61.63; H, 6.71; N, 7.18. Found: C, 61.42; H, 6.85; N, 7.23.

^1^H NMR (600 MHz, CDCl_3_) δ 8.79 (d, 1H, *J* = 1.68 Hz), 8.47 (d, 1H, *J* = 2.82 Hz), 7.79 (dd, 1H, *J* = 1.68 Hz), 4.38 (t, 2H, *J* = 6.60 Hz), 1.79 (quin, 2H, *J* = 6.66 Hz), 1.60 (s, br, 1H), 1.50 (hex, 2H, *J =* 7.38 Hz), 1.01 (t, 3H, *J =* 7.38 Hz).

^13^C NMR, proton decoupled (150.9 MHz, CDCl_3_) δ 165.1, 153.9, 141.5, 140.7, 127.9, 124.7, 65.6, 30.6, 19.2, 13.7.

***n*-Butyl 5-(prop-2-yn-1-yloxy)nicotinate** (**2**). To a solution of **1** (481.8 mg, 2.48 mmol) in anhydrous dimethylformamide (10 mL), cesium carbonate (1.2 g, 3.72 mmol, 1.5 equiv.) was added in one portion under an inert atmosphere. Subsequently, propargyl bromide (80 wt.% solution in toluene, 553.3 mg, 0.63 mmol, 0.4 mL, 1.5 equiv.) was added dropwise to the reaction mixture. The reaction was stirred at room temperature for 1 hour and monitored by thin-layer chromatography (TLC) (30% ethyl acetate-hexane). Upon completion, as indicated by the disappearance of the starting material, the reaction was quenched by the addition of 1 N HCl until the pH reached approximately 6. The mixture was frozen and lyophilized overnight. The resulting solid was dissolved in ethyl acetate and filtered to remove insoluble impurities. The filtrate was then purified via flash column chromatography on silica gel using ethyl acetate-hexane as the eluent. Two fractions were collected, with the second fraction containing the pure compound **2**. The purified fraction was concentrated under reduced pressure, yielding a slightly yellow solid (0.54 g, 93.75%).

Characterization: m.p. 44–48 °C reported. m.p. 45–47 °C [5]); TLC R_f_ = 0.54 (30% ethyl acetate-hexane).

An analytical sample was prepared by dissolving a small portion (100 mg) in hot methanol (0.25 mL), and adding hot water (ca. 1 mL) followed by rapid cooling. Once crystals began to form an additional 5 mL of water was added, the solids collected by centrifugation and washed with water. White crystals (89 mg) were obtained: m.p. 46-48 °C.

Anal. Calcd. for C_13_H_15_NO_3_: C, 66.94; H, 6.48; N, 6.00. Found: C, 66.83; H, 6.56; N, 6.06.

^1^H NMR (600 MHz, CDCl_3_) δ 8.87 (d, 1H, *J* = 1.62 Hz), 8.53 (d, 1H, *J =* 3.00 Hz), 7.88 (dd, 1H, *J* = 1.68 Hz), 4.81 (d, 2H, *J* = 2.40 Hz), 4.36 (t, 2H, *J* = 6.66 Hz), 2.64 (t, 1H, *J* = 2.40 Hz), 1.77 (quin, 3H, *J* = 6.66 Hz), 1.48 (hex, 2H, *J =*7.38 Hz), 0.98 (t, 3H, *J =* 7.44 Hz);

^13^C NMR, proton decoupled (150.9 MHz, CDCl_3_) δ 165.1, 153.4, 143.7, 142.7, 126.8, 121.6, 77.1, 76.9, 65.4, 56.3, 30.6, 19.2, 13.7.

**5-(Prop-2-yn-1-yloxy)nicotinic acid** (**3**). Compound **2** (129.9 mg, 0.56 mmol) was dissolved in 3 mL of methyl alcohol. Aqueous 4 N NaOH (2 mL) was added and stirred at room temperature for 2 hr. Once the TLC showed consumption of all the starting material, 8 mL of 1 N HCl was added and stirred at room temperature for 30 min. After 30 min of stirring, the pH was tested to ensure the pH was neutral or slightly acidic. The solvent was evaporated *in vacuo*, and to the dried residue was added 30 mL of ethyl acetate and transferred to a separatory funnel. The organic layer was washed twice with 10 mL of water. The ethyl acetate extracts were dried over Na_2_SO_4_ and concentrated by distillation *in vacuo*. The crude product was further purified by recrystallizing from a large amount of water to give white solid crystals. (80 mg, 81%). m.p. 205-207 °C, (reported m.p. 205-207 °C [5]); TLC R_f_ 0.13 (50% hexane/ethyl acetate).

^1^H NMR (600 MHz, CD_3_OD) δ 8.77 (d, 1H, *J* = 1.32 Hz), 8.50 (d, 1H, *J* = 2.82 Hz), 8.02 (q, 1H *J* = 1.62 Hz), 4.93 (s, 2H), 3.10 (t, 1H, *J* = 2.34 Hz);

^13^C NMR, proton decoupled (150.9 Hz, D_2_O) δ 166.2, 154.1, 142.6, 141.5, 127.7, 122.4, 77.1, 76.8, 55.9.

Anal. Calcd. for C_9_H_7_NO_3_: C, 61.02; H, 3.98; N, 7.91. Found: C, 60.90; H, 4.37; N, 7.62.

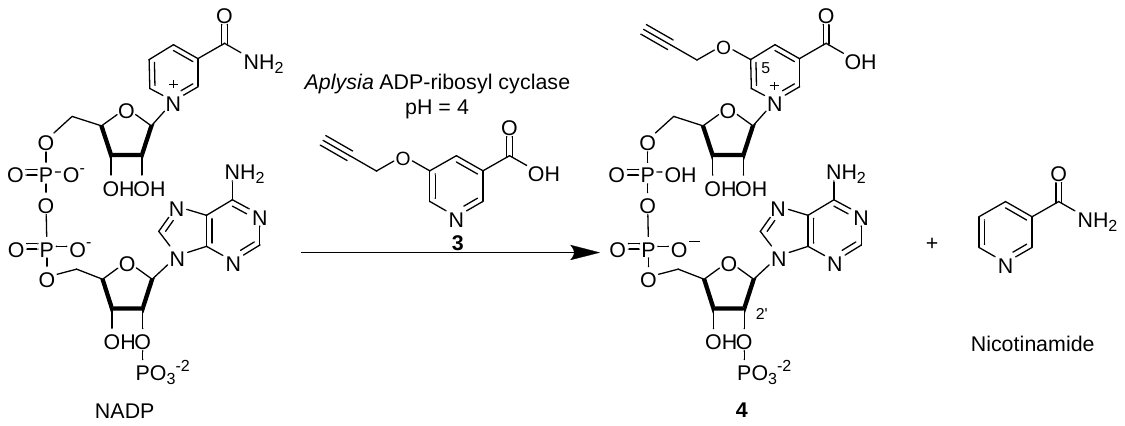

**Scheme S2.** Enzyme catalyzed pyridine base exchange reaction for the synthesis of 5-(prop-2-yn-1-yloxy)-NAADP (**4**) from NADP and 5-(Prop-2-yn-1-yloxy) nicotinic acid (**3**)

**Synthesis of** **5-(prop-2-yn-1-yloxy)-NAADP** (**4**). Nicotinic acid derivative **3** (53.1 mg, 0.3 mmol) was dissolved in dimethyl sulfoxide (2 mL) and distilled water (6 mL) and transferred to a 20 mL vial that was equipped with a stirring bar. The reaction solution was continuously stirred for 30 min in an incubator (37 °C). After this time, the pH of the reacting solution was adjusted to about 5 using 0.1-0.5 M NaOH until most of compound **3** was in solution. NADP (15 mg, 0.015mmol) was added to the reaction solution and stirred at 37 °C for another 10 min and the pH readjusted to about 5. Next, wild-type *Aplysia californica* ADP-ribosyl cyclase [2](360 μL, 72 μg) was added to the reaction and stirred at 37 °C for 3 hours. 5-(Prop-2-yn-1-yloxy)-NAADP was purified by anion exchange chromatography on AG MP-1 using a linear gradient formed between water and 150 mM trifluoroacetic acid. The fractions containing **4** were combined, the pH was adjusted to 7 using 0.1 M NaOH, and the solution was then loaded on a DEAE cellulose anion-exchange column and the chromatography developed by applying a linear gradient of 0-400 mM NH_4_HCO_3_ over a total volume of 1.2 L. The product eluted into ca. 120-180 mM NH_4_HCO_3_. The fractions displayed a UV absorbance at 260 nm were combined, frozen at -80 °C, and lyophilized to afford a white powder. The lyophilized dry powder was re-dissolved in 5 mL of water and lyophilized again (3×) until salt free 5-(prop-2-yn-1-yloxy)-NAADP was obtained as an off-white powder (11.8 mg, 73.3 %). UV/Vis (H_2_O) λ_max_ 258 nm (ε_259_ = 14,600 L mol^-1^ cm^-1^) with a shoulder at 294 nm (A_258_/A_294_ = 2.62). See Figure S1.

^1^H-NMR (600 MHz, D_2_O) δ 8.79 (s, 1H) 8.64 (s, 1H), 8.44 (s, 1H), 8.41 (s, 1H), 8.15 (s, 1H), 6.14 (d, 1H, J = 5.7), 5.97 (d, 1H, J = 3.9), 5.04 (m, 1H), 4.99 (s, 2H), 4.62-4.18 (m, 9H), 3.06 (s, 1H).

^13^C-NMR, proton decoupled (150.903 MHz, D_2_O) δ 166.7, 156.7, 153.5, 150.1, 148.7, 140.8, 137.7, 134.5,132.0, 128.9, 118.2, 99.8, 86.5 (d, J_P-C_ = 7.5 Hz), 85.9 (d, J_P-C_ = 6 Hz), 83.5 (d, J_P-C_ = 9 Hz), 78.8, 77.3, 76.6 (d, J_P-C_ = 4.5 Hz), 76.2, 70.3, 69.8 (d, J_P-C_ = 3 Hz), 65.2 (d, J_P-C_ = 4.5 Hz), 64.9 (d, J_P-C_ = 4.5 Hz), 57.9.

**Synthesis of BODIPY-(EG_4_)-NAADP (5).** The strategy for this synthesis is to produce a BODIPY FL coupled through an amide linkage to a tetra-ethylene glycol spacer that terminates in an azide. The BODIPY-EG_4_-N_3_ azide can then be coupled to NAADP analog **4** using a click reaction.

Step1. Activation of BODIPY as the NHS-ester.

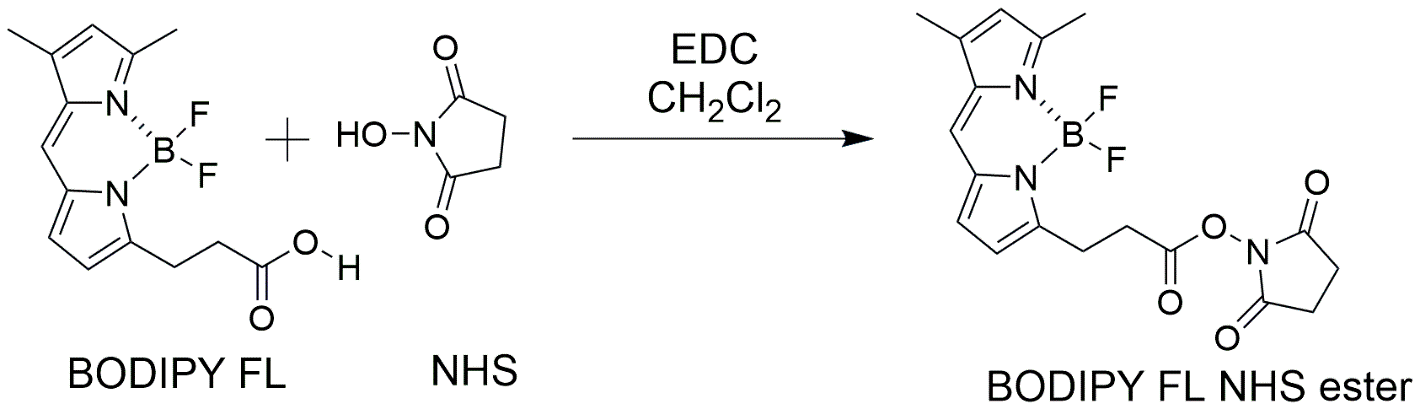

Scheme S3. Synthesis of BODIPY FL N-hydroxy succinimide ester.

The method is to treat BODIPY carboxylic acid with an excess of *N*-hydroxysuccinimide (NHS) and 1-ethyl-3-(3-dimethylaminopropyl)carbodiimide hydrochloride (EDC⦁HCl) as the coupling agent in methylene chloride to form the BODIPY-NHS active ester. Excess NHS, EDC⦁HCl, and the coproduct 1-ethyl-3-(3-dimethylaminopropyl)urea hydrochloride are completely removed by extraction with water, the methylene chloride was dried above anhydrous magnesium sulfate, filtered, and the solvent evaporated to leave the pure BODIPY-NHS.

Procedure 1. BODIPY FL (102 mg; 0.349 mmol) was dissolved in 15 mL of methylene chloride, 2 eq of *N*-hydroxysuccinimide (80.33 mg; 0.698 mmol; 2-fold excess) was added followed by EDC⦁HCl (133.8 mg; 0.698 mmol; 2-fold excess). The reaction mixture was stirred at room temperature overnight. The methylene chloride solution was washed twice with 50 mL portions of water to remove the carbodiimide, the urea co-product, and excess NHS. The organic phase was dried over anhydrous magnesium sulfate, filtered (fluted filter paper), and the solvent removed under reduced pressure to leave colored residue. TLC: R*_f_* =0.8 (5% MeOH in DCM). If desired the mixture could be purified quickly by automated flash column chromatography on silica gel (40 g silica, 5% MeOH in DCM) to provide compound BODIPY FL NHS ester as a red solid (82.6 mg, 68.3%). UV/Vis (methanol) λ_max_ 504 nm.

^1^H-NMR (600 MHz, CDCl_3_) δ 7.10 (s, 1H), 6.89 (d, 1H, J = 3.96), 6.34 (d, 1H, J = 3.96), 6.13 (s, 1H), 3.39 (t, 2H, J = 7.32), 3.09 (t, 2H, J = 7.38), 2.84 (br, 4H), 2.57 (s, 3H), 2.55 (s, 3H).

Step 2. Preparation of BODIPY-EG_4_-N_3_.

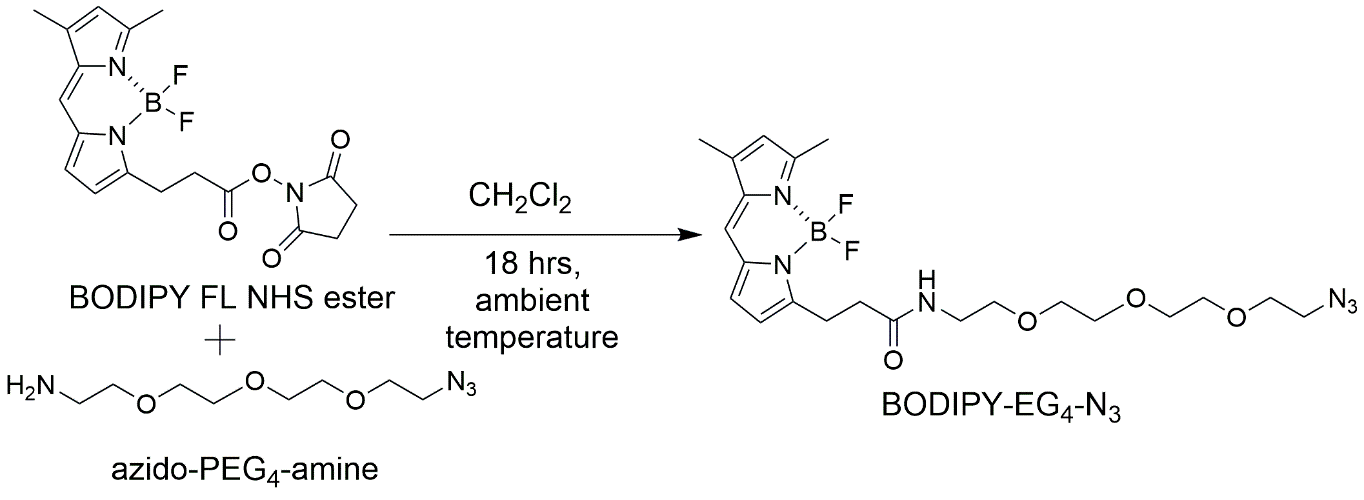

Scheme S4. Preparation of the BODIPY-EG_4_-Azide

The method is to treat a concentrated solution of the BODIPY FL active ester with the azido-PEG_4_-amine in methylene chloride followed by chromatographic purification.

Procedure 2. One equivalent of BODIPY FL NHS ester (100 mg, 0.26 mmol) and 2 equivalent of 11-azido-3,6,9-trioxaundecan-1-amine (azido-PEG_4_-amine) (113.5 mg, 0.52 mmol, 103.2 µl) was dissolved in 0.5 ml of DCM and mixed at room temperature for 18 hr. After 18 hr, the reaction mixture was diluted with DCM, loaded on a silica gel column, and further purified by automated flash column chromatography on silica gel.

^1^H-NMR (600 MHz, CDCl_3_) δ 7.27 (s, 1H), 7.09 (s, 1H), 6.88 (d, 1H, J = 3.96), 6.30 (d, 1H, J = 3.96), 6.21 (br, 1H), 6.12 (s, 1H), 3.69-3.35 (m, 16H), 3.28 (t, 2H, J = 7.5), 2.63 (t, 2H, J = 7.68), 2.56 (s, 3H), 2.25 (s, 3H).

^13^C-NMR, proton decoupled (150.903 MHz, CDCl_3_) δ 170.7, 159.0, 156.8, 142.7, 134.0, 132.4, 127.3, 122.7, 119.3, 116.5, 69.6, 69.2, 69.0, 68.8, 49.6, 38.2, 34.9, 23.8, 13.9, 10.2.

Step 3. Click reaction between BODIPY- EG_4_-N_3_ and 5-(prop-2-yn-1-yloxy)-NAADP (**4**) resulting in BODIPY-(EG_4_)-NAADP (**5**)
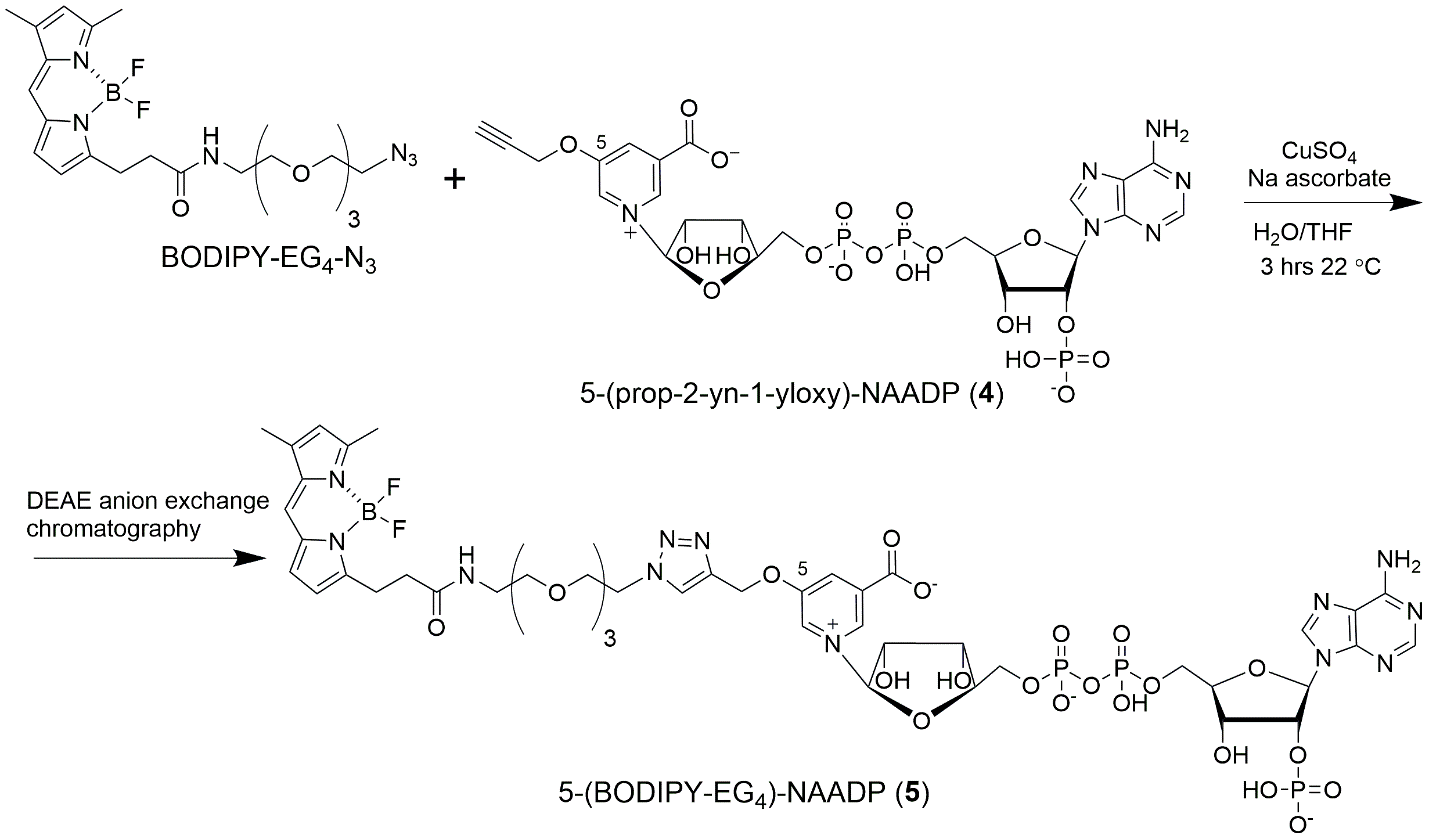

Scheme S5. Cu-catalyzed click synthesis of BODIPY-(EG_4_)-NAADP (**5**).

**Synthesis of BODIPY-(EG_4_)-NAADP (5).** The method is to mix a solution of BODIPY-EG_4_-N_3_ dissolved in 3:1 tetrahydrofuran (THF)-water with a concentrated 50 mM aqueous solution of 5-(propargyl ether)-NAADP (**4**) and initiate a “click” reaction connecting the azide to the alkyne as a triazole. The click reaction is initiated by adding a 10% aqueous solution of CuSO_4_⦁6H_2_O followed by a 100 mg/mL solution of sodium ascorbate. The click reaction is over within 3 hours and the product is purified by anion exchange chromatography on DEAE Sephacel.

Procedure 3. An approximately 50 mM solution of the clickable 5-(prop-2-yn-1-yloxy)-NAADP (**4**), ammonium salt form, in H_2_O was prepared by dissolving 10.2 mg (12 µmol; 1 equivalent) of the dinucleotide in 240 µL of water. The solution was placed in a 1.5 mL Eppendorf tube and BODIPY-EG_4_-N_3_ (11.8 mg; 24 µmol; 2 equivalents) was added as a solution in 480 µL of 3:1 THF-H_2_O. The solutions were mixed and all liquid returned to the bottom of the tube by a brief centrifugation. Next, 192 µL of a 10% (w/v) solution of copper (II) sulfate hexahydrate (1.92 mg; 7.7 µmol; 0.6 equivalent) followed by 143 µl of freshly prepared 100 mg/mL sodium ascorbate in dI⦁H_2_O (14.3 mg; 72.2 µmol, 6 equivalents) were added sequentially to the reaction. Mixing and centrifugation was repeated to concentrate the materials at the bottom of the Eppendorf tube, the reaction incubated at ambient temperature for 3 hours and at its conclusion diluted with 10 mL of dI⦁H_2_O. The reaction mixture was loaded onto a DEAE-anion exchange column (1.5 cm × 15 cm, 26.5 mL bed volume) that had been equilibrated with 10 mM ammonium bicarbonate (NH_4_HCO_3_) at pH 7.5. The chromatography was developed by applying a linear gradient formed between 600 mL of 0.010 M NH_4_HCO_3_ and 600 mL of 0.4 M NH_4_HCO_3_. The flow rate was ca. 2 mL /min, 10 mL fractions were collected and the absorbance at 260 nm and 503 nm of effluent fractions was determined. The product eluted into fractions 64, (ca. 210 mM NH_4_­HCO_3_), through 74, (about 247 mM NH_4_­HCO_3_). The fractions displayed a UV absorbance at both 503 nm and 260 nm were collected, combined, frozen, and lyophilized to afford a red powder. The lyophilized dry powder was re-dissolved in 5 mL of water and lyophilized again (3×) until a pure salt free compound was obtained as a red powder (9.6 mg, 61.9 %). UV/Vis (H_2_O) λ_max_ 259 nm (ε_259_ = 15,400 L mol^-1^ cm^-1^) 294 nm, and 506 nm (ε_506_ = 42,000 L mol^-1^ cm^-1^). See figure S2.

^1^H-NMR (600 MHz, D_2_O) δ 8.69 (s, 1H), 8.53 (s, 1H), 8.39 (s, 1H), 8.27 (s, 1H), 8.18 (s, 1H), 8.01 (s, 1H), 7.07 (s, 1H), 6.85 (d, 1H, J = 3.2), 6.22 (d, 1H, J = 3.5), 6.13 (s, 1H), 6.00 (d, 1H, J = 4.5), 5.84 (d, 1H, J = 5.3), 5.25 (dd, 2H, J_1_ = 11.9, J_2_ = 52.6), 4.58-3.91 (m, 9H), 3.56-3.32 (m, 16 H), 3.05 (t, 2H, J = 6.4), 2.59 (t, 2H, J = 7.0), 2.35 (s, 3H), 2.12 (s, 3H).

^13^C-NMR, proton decoupled (150.903 MHz, D_2_O) δ 174.9, 161.3, 157.1, 155.6, 154.5, 151.6, 146.0, 134.9, 134.2, 132.8, 132.1, 128.6, 127.9, 124.2, 121.0, 116.5, 99.7, 86.6, 85.8, 83.4, 77.4, 76.6, 70.3, 69.9, 69.64, 69.58, 69.43, 69.38, 68.8, 68.7, 65.2, 64.9, 62.9, 50.1, 39.1, 34.3, 30.3, 26.6, 24.3, 24.1, 23.8, 14.1, 12.7, 10.5.

Positive mode MALDI mass spectra: Calcd. m/z for C_46_H_61_BF_2_N_12_O_23_P_3_, [M]^+^ 1291.3241; measured m/z was 1291.169 (120 ppm error).

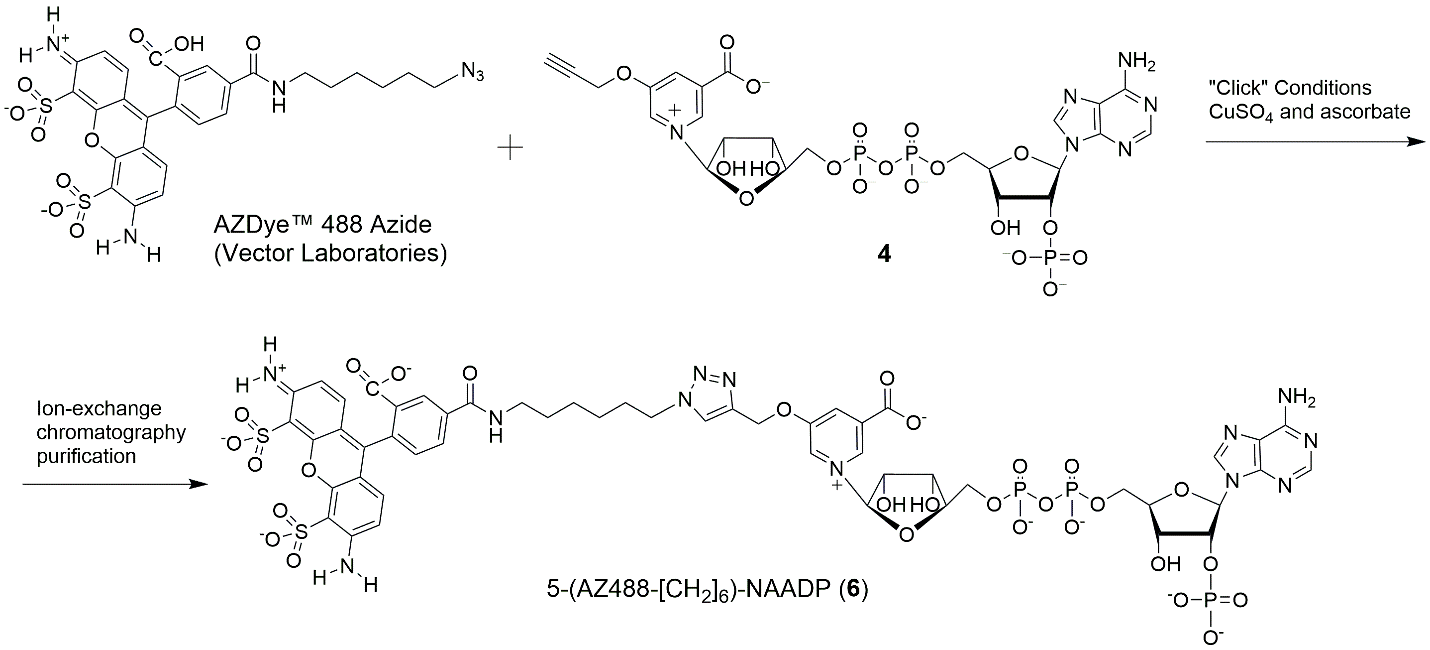

Scheme S6**.** Cu-catalyzed click synthesis of 5-(AZ488-[CH_2_]_6_)-NAADP (**6**).

**Synthesis of AZ488 with hydrophobic spacer arm (6)** AZdye488^®^ azide (5 mg, 0.0076 mmol; 1 equivalent; Vector labs, Newark, CA, USA; #CCT-1275-5) was dissolved in 76 μL H_2_O and 76 μL THF to make 152 µL of a 50 mM solution and mixed with 228 μL of a 50 mM solution of clickable 5-(prop-2-yn-1-yloxy)-NAADP (**4**) in water ( 9.7 mg, 0.011 mmol, 1.5 equivalent), 12.5 μl CuSO_4_⦁6H_2_O solution (10 mg/ml in ddH_2_O) (0.1 equivalent), and 31 μL of freshly prepared sodium ascorbate solution (100 mg/ml in ddH_2_O) (2 equivalents). The reaction mixture was incubated at 0 °C to 25 °C for 2 hr. After 2 hr, the reaction mixture was diluted with 10 mL ddH_2_O and applied to a DEAE cellulose anion-exchange column (1.5 cm × 15 cm, 26.5 mL bed volume) that had been equilibrated with 10 mM ammonium bicarbonate (NH_4_HCO_3_) at pH 7.5. The chromatography was developed by applying a linear gradient formed between 600 mL of 0.010 M NH_4_HCO_3_ and 600 mL of 0.4 M NH_4_HCO_3_. The flow rate was ca. 2 mL /min, 10 mL fractions were collected and the absorbance at 260 nm and 494 nm of effluent fractions was determined. The product eluted at about fraction 59, at a salt concentration of approximately 295 mM NH_4_HCO_3_, and ended at fraction 72, ca. 360 mM NH_4_HCO_3_. The fractions displaying a UV absorbance at both 260 nm, and 494 nm were combined, frozen, and lyophilized to afford a purple powder. The lyophilized dry powder was re-dissolved in 5 mL of water and lyophilized again (3×) until a pure compound **6** was obtained orange powder (6.9 mg, 62.7 %). UV/Vis (H_2_O) showed absorption maxima (λ_max_) at 241 nm (ε_241_ = 53,400 L mol^-1^ cm^-1^), 305 nm, and 493 nm (ε_494_ = 70,500 L mol^-1^ cm^-1^). See Figure S3.

^1^H-NMR (600 MHz, D_2_O) δ 8.82 (s, 1H), 8.70 (s, 1H), 8.45 (s, 1H), 8.38 (s, 1H), 8.19 (s, 1H), 8.11 (s, 1H), 8.02(d, 1H, J = 8.10), 7.98 (d, 1H, J=8.10), 7.66 (s, 1H), 7.14 (q, 2H, J = 8.66), 6.85 (q, 2H, J = 9.06), 6.08( d, 1H, J = 5.52), 6.01 (d, 1H, J = 4.92), 5.37 (dd, 2H, J_1_ = 11.8, J_2_ = 38.5), 5.02 (m, 1H), 4.56-4.10 (m, 9H), 3.31 (t, 2H, J = 6.3),1.83-1.17 (m, 10H).

^13^C-NMR, proton decoupled (150.903 MHz, D_2_O) δ 173.0, 169.1, 166.7, 161.0, 157.2, 155.5, 155.2, 153.5, 149.0, 141.9, 141.3, 140.0, 137.9, 135.2, 134.5, 132.4, 131.9, 131.8, 131.4, 129.4, 128.8, 128.4, 127.9, 125.4, 119.5, 118.4, 113.6, 111.1, 99.8, 86.7, 86.0 (d, J = 8.6 Hz), 83.0 (d, J = 7.9 Hz), 77.4, 75.8, 70.4, 70.0, 65.3, 65.0, 63.0, 50.2, 39.8, 29.0, 28.0, 25.2, 25.0.

Negative mode MALDI mass spectra: Calcd. m/z for (C_51_H_54_N_12_O_29_P_3_S_2_)^-1^, m/z 1455.1779; measured m/z: 1455.1731 (3.3 ppm error).

**Figures to Accompany Supplementary Material**

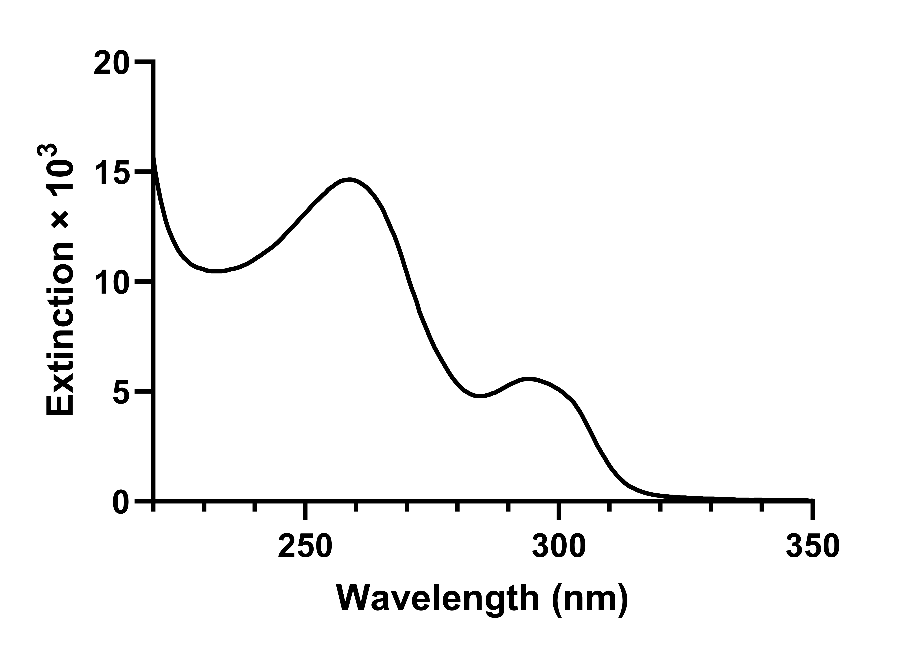

Figure S1. Electronic absorption spectrum of 5-(prop-2-yn-1-yloxy)-NAADP (**4**).

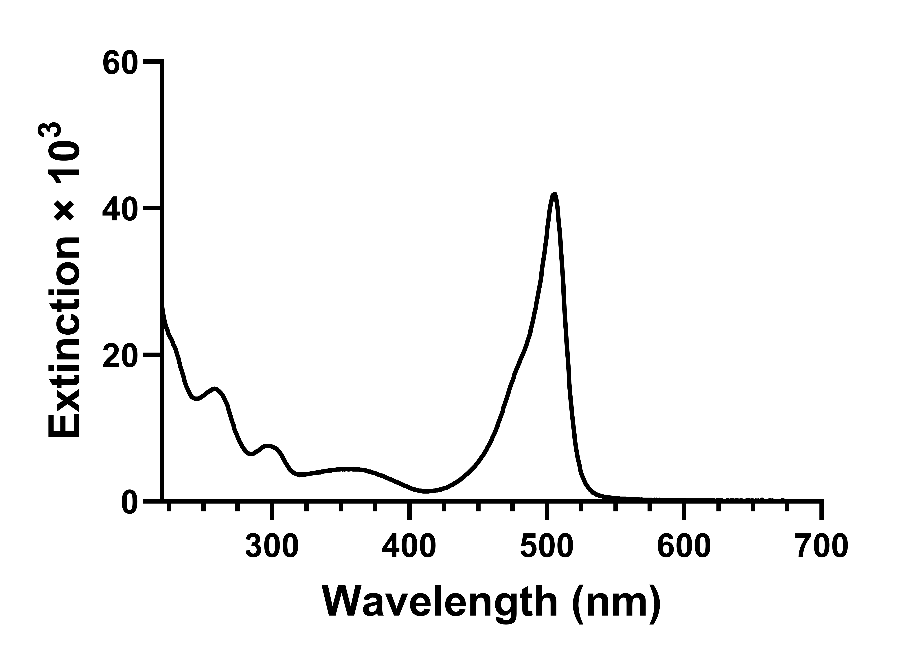

Figure S2. Electronic absorption spectrum of BODIPY-(EG_4_)-NAADP (**5**).

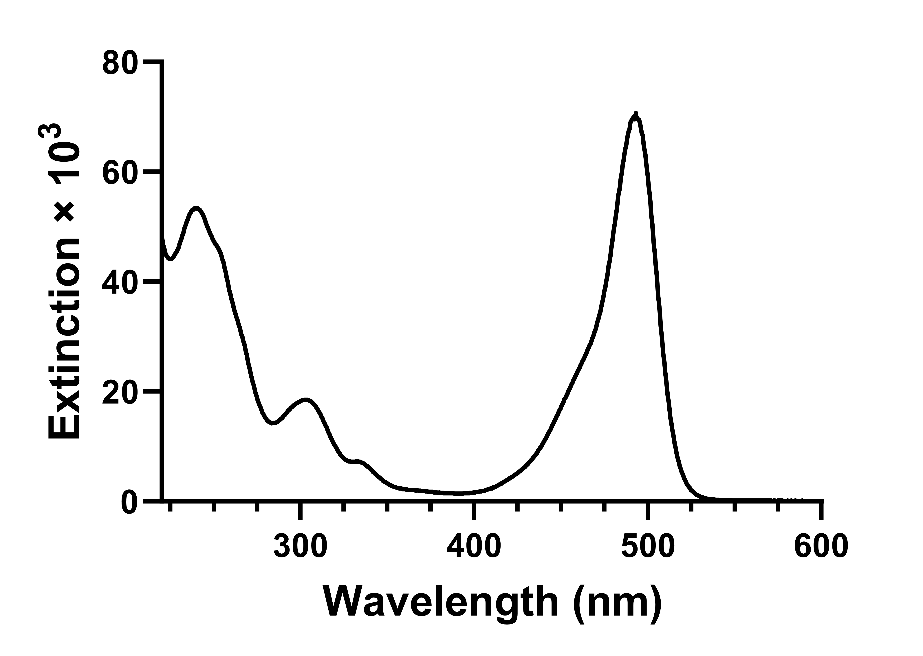

Figure S3. Electronic absorption spectrum of 5-(AZ488-[CH_2_]_6_)-NAADP (**6**).

**Supplementary Methods References**

1.	Serdjukow, S.; Kink, F.; Steigenberger, B.; Tomás-Gamasa, M.; Carell, T., Synthesis of γ-labeled nucleoside 5′-triphosphates using click chemistry. *Chem Commun* **2014,** 50, (15), 1861-1863.

2.	Munshi, C.; Lee, H. C., High-Level Expression of Recombinant *Aplysia* ADP-Ribosyl Cyclase in *Pichia pastoris* by Fermentation. *Protein Expr Purif* **1997,** 11, (1), 104-110.

3.	Jain, P.; Slama, J. T.; Perez-Haddock, L. A.; Walseth, T. F., Nicotinic acid adenine dinucleotide phosphate analogues containing substituted nicotinic acid: effect of modification on Ca^(2+)^ release. *J Med Chem* **2010,** 53, (21), 7599-612.

4.	Lee, H. C.; Aarhus, R., Structural determinants of nicotinic acid adenine dinucleotide phosphate important for its calcium-mobilizing activity. *J Biol Chem* **1997,** 272, (33), 20378-83.

5.	Asfaha, T. Y. Clickable, Photoactive NAADP Analogs for Isolation and Purification of the Unknown NAADP Receptor. Doctoral Dissertation, University of Toledo, Toledo Ohio USA, 2016.

6.	Alexson, J. T.; Bodley, J. W.; Walseth, T. F., A volatile liquid chromatography system for nucleotides. *Analytical Biochemistry* **1981,** 116, 347-360.

7.	Guan, Z.; Slama, J. T., Synthesis of a Stable Long-Wavelength Fluorescent BODIPY FL-NAADP Conjugate. *Molbank* **2025,** 2025, (4), M2085.
